# Development of motor neurons and motor activity in zebrafish requires F-actin nucleation by Fmn2b

**DOI:** 10.1101/2021.08.10.455777

**Authors:** Dhriti Nagar, Blake Carrington, Shawn M Burgess, Aurnab Ghose

**Author notes:** Correspondence: Dr. Aurnab Ghose, Indian Institute of Science Education and Research (IISER) Pune, Dr Homi Bhabha Road, Pune 411008, INDIA.

## Abstract

**Background:** Cytoskeletal remodelling plays a pivotal role in the establishment of neuronal connectivity during development and in plasticity in adults. Mutations in the cytoskeleton regulatory protein Formin-2 (Fmn2) are associated with neurodevelopmental disorders like intellectual disability, though its function in neuronal morphogenesis has not been characterised *in vivo*.

**Results:** Here we develop a loss-of-function model for *fmn2b*, the zebrafish orthologue of Fmn2, using CRISPR/Cas9-mediated gene editing. *fmn2b* mutants display motor deficits starting from the earliest motor responses in the embryo. We find that *fmn2b* is expressed in spinal motor neurons and its loss reduces motor neuron innervation of the axial muscles without affecting myotome integrity. The translocation of caudal primary (CaP) motor neuron outgrowth is compromised in *fmn2b* mutants, while rostral primary (RoP) motor neurons have missing soma or stall at the horizontal myoseptum. Strikingly, axon collateral branching of the motor neurons is severely compromised and results in reduced synaptic coverage of the myotome. Rescue experiments identify the requirement for Fmn2-mediated actin nucleation for motor neuron outgrowth and arborisation.

**Conclusions:** The zebrafish loss-of-function model of Fmn2 reveals the specific requirement of F-actin polymerisation by Fmn2 in neuromuscular development. It also underscores the role of Fmn2 in motor neuropathies, especially as a proportion of individuals harbouring mutations in Fmn2 present with hypotonia.

## INTRODUCTION

Developing neurons migrate to their predetermined targets, undergo axonogenesis, pathfinding and eventually arborize to make functional synapses. Cytoskeletal remodelling allows dynamic interactions of the developing neurons with the environment during outgrowth, pathfinding and arborization (Dickson, 2002; Lowery and Van Vactor, 2009; Gordon-Weeks and Fournier, 2014; Flynn and Bradke, 2020) aided by several actin remodelling proteins (Dent et al., 2011; Kessels et al., 2011; Lewis et al., 2013; Coles and Bradke, 2015; Armijo-Weingart and Gallo, 2017).

Remodelling of actin filaments allows protrusive structures to probe the environment for cues in the form of growth cone filopodia and axonal and dendritic branches (Lowery and Van Vactor, 2009; Lewis et al., 2013; Armijo-Weingart and Gallo, 2017; Menon and Gupton, 2018).

In zebrafish, spinal motor neurons exit perpendicular to the spinal cord and traverse the myotomes to arborize and make synapses onto the axial muscles (Myers et al., 1986; Westerfield et al., 1986). Adequate innervation of muscles by the motor neurons is ensured by hemisegment specific outgrowth, pathfinding and arborization. Precise neuronal morphogenesis of motor neurons is critical for achieving context-specific synaptic drive at the level of muscles, enabling the organism to execute motor behaviours.

Actin nucleating proteins are a major class of actin remodelling factors and have been implicated in neuronal morphogenesis. The actin nucleator Cobl localizes to regions with high actin dynamics and is implicated in the maintenance of neurite branching (Ahuja et al., 2007). Actin nucleators mediate the local rearrangement of actin filaments via the formation of dynamic structures like F-actin patches and F-actin blobs and regulate the initiation of protrusions and branching (Ketschek and Gallo, 2010; Spillane et al., 2011; Spillane and Gallo, 2014; Nithianandam and Chien, 2018; Kundu et al., 2020). Reports have characterized the role of Dishevelled associated activator of morphogenesis (Daam) formins in *Drosophila* axonal growth (Matusek et al., 2008; Prokop et al., 2011) and Daam1a in habenular morphogenesis and IPN connectivity in zebrafish (Colombo et al., 2013). Another study identified Formin-3 (Form3) in the maintenance of complex dendritic arbours of nociceptive sensory neurons in *Drosophila* via microtubule stabilization and adequate organelle trafficking (Das et al., 2021). However, the function of formin family of actin nucleators in axonal branching *in vivo* remains elusive.

Formin-2 (Fmn2) is part of the FMN family of formins and is known to be involved in neuronal development. Biallelic mutations in Fmn2 cause neurodevelopmental defects resulting in intellectual disability and muscle hypotonia in humans (Law et al., 2014). Primary hippocampal neurons from Fmn2 null mice show defective dendritic spine morphogenesis (Law et al., 2014) and there is increasing evidence implicating copy number variations (CNVs) and single nucleotide polymorphisms (SNPs) in Fmn2 in sporadic Amyotrophic Lateral Sclerosis (SALS) (Schymick et al., 2007; Pamphlett et al., 2011; Morello et al., 2018). Fmn2 has previously been characterized in outgrowth, pathfinding (Sahasrabudhe et al., 2016; Ghate et al., 2020; Kundu et al., 2021) and axonal branching (Kundu et al., 2020) in chick spinal primary neurons *in vitro*. Depletion of Fmn2 causes reduced growth cone motility and mechanotransduction in chick spinal primary neurons (Sahasrabudhe et al., 2016; Ghate et al., 2020). Fmn2 regulates the stabilization of actin patches required for the initiation of axonal branching in cultured chick spinal neurons by insulating the actin patch from actin depolymerizing factors like ADF (Kundu et al., 2020). Morpholino-mediated transient knockdown of Fmn2b, the zebrafish ortholog of Fmn2, causes impaired outgrowth of the spiral fiber neuron commissures in the hindbrain and results in diminished short latency escape response (Nagar et al., 2021).

In this study, we generate *fmn2b* mutants using the CRISPR-Cas9 based gene editing to assess the contribution of Fmn2b in neural circuit development and behaviour and to improve our understanding of the mechanistic underpinnings of the neurodevelopmental disorders associated with Fmn2.

Motor neurons are the final, executory neurons driving motor responses. In zebrafish, the spinal motor neurons receive synaptic inputs from reticulospinal neurons, including the Mauthner cell and several hindbrain interneurons, which regulate motor neuron function in response to stimuli (Bernhardt et al., 1990; Hale et al., 2001; McLean and Fetcho, 2008). Three subtypes of primary motor neurons are present on either side of the spinal cord – the Caudal primary (CaP), the Middle primary (MiP) and the Rostral primary (RoP) – with distinct morphology and muscle targets (Eisen et al., 1986; Myers et al., 1986; Bernhardt et al., 1990; Melançon et al., 1997; Stifani, 2014). The myotomes are also innervated by secondary motor neurons which develop after the primary motor neurons and have less complex arborization fields (Pike et al., 1992; Beattie et al., 1997; Menelaou and Svoboda, 2009). Motor neurons innervate the target muscles to form structurally and functionally distinct motor units and regulate the overall synaptic drive of the muscles during motor behaviours (Saint-Amant and Drapeau, 1998; Drapeau et al., 2002; Bagnall and McLean, 2014; Bello-Rojas et al., 2019).

This study uncovers the role of the zebrafish orthologue of Formin-2, Fmn2b, in regulating spinal motor neuron outgrowth and branching *in vivo* and the consequent effects on motor behaviour.

## RESULTS

### Generating fmn2b CRISPR mutants

Zebrafish has two paralogs of Formin-2, *fmn2a* and *fmn2b*. *fmn2b*, located on chromosome 12, has higher sequence homology with human Fmn2 compared to *fmn2a* and has the characteristic Formin Homology 1 (FH1), Formin Homology 2 (FH2) and Formin Spire Interaction (FSI) domains (Nagar et al., 2021). In contrast, the FH1 domain of *fmn2a* is truncated. As observed in humans, rodents and chicks (Leader and Leder, 2000; Sahasrabudhe et al., 2016), *fmn2b* mRNA is enriched in the zebrafish nervous system whereas *fmn2a* shows no detectable expression in the nervous system of zebrafish (Nagar et al., 2021). In this study, we used the CRISPR-Cas9 technology to generate mutants of *fmn2b*, the functional ortholog of mammalian Fmn2 in zebrafish.

To generate *fmn2b* knockout zebrafish, sgRNAs targeting the exon 1 of *fmn2b* were designed using the <u>CRISPRscan</u> tool (Moreno-Mateos et al., 2015) and cross verified for high scores using the <u>ZebrafishGenomics</u> track (LaFave et al., 2014) in UCSC genome browser. Exon 1 was targeted to ensure that the resulting mutant would lack the functional domains.

The embryos injected with sgRNA and Cas9 mRNA were raised till sexual maturity. Individual embryos from the injected clutch were outcrossed with wildtype fish of the opposite sex to obtain eggs that were genotyped using fluorescent PCR (Varshney et al., 2016). Multiple founder lines with different allelic mutations were identified for *fmn2b* by fluorescent-PCR. The F1 progeny obtained from the outcross of these founder lines was raised to adulthood and the siblings were in-crossed. The F2 progeny were raised to adulthood and genotyped using the caudal fin clip method and subsequent Sanger sequencing of the genomic locus flanking the sgRNA-1 target sequence to determine whether they were heterozygous or homozygous for the inherited mutant allele.

The sgRNA caused various indels near the intended target around 696 bp into the *fmn2b* cDNA sequence in exon 1, corresponding to 232^nd^ amino acid position in the 1454 amino acid long Fmn2b protein sequence. Two alleles causing a 4 bp and a 7 bp deletion, respectively, were selected to obtain homozygous mutants *fmn2b^Δ4/Δ4^* and *fmn2b^Δ7/Δ7^* (Figure 1 A-B). The 4 bp deletion in the *fmn2b^Δ4/Δ4^* mutant leads to frameshift variant (p.Leu233TrpfsTer281) with Tryptophan as the first amino acid changed in place of Leucine at 233^rd^ amino acid position causing premature stop codon at the 281^st^ amino acid. The 7 bp deletion acid in the *fmn2b^Δ7/Δ7^* results in the frameshift variation (p.Val234ThrfsTer280) with Threonine replacing Valine as the 234^th^ amino acid and causing a premature stop codon at 280^th^ amino acid position (Figure 1 C). The two homozygous mutant alleles were in-crossed to obtain heteroallelic *fmn2b* mutants (*fmn2b^Δ4/Δ7^*) and used in subsequent experiments.

**Figure 1.**
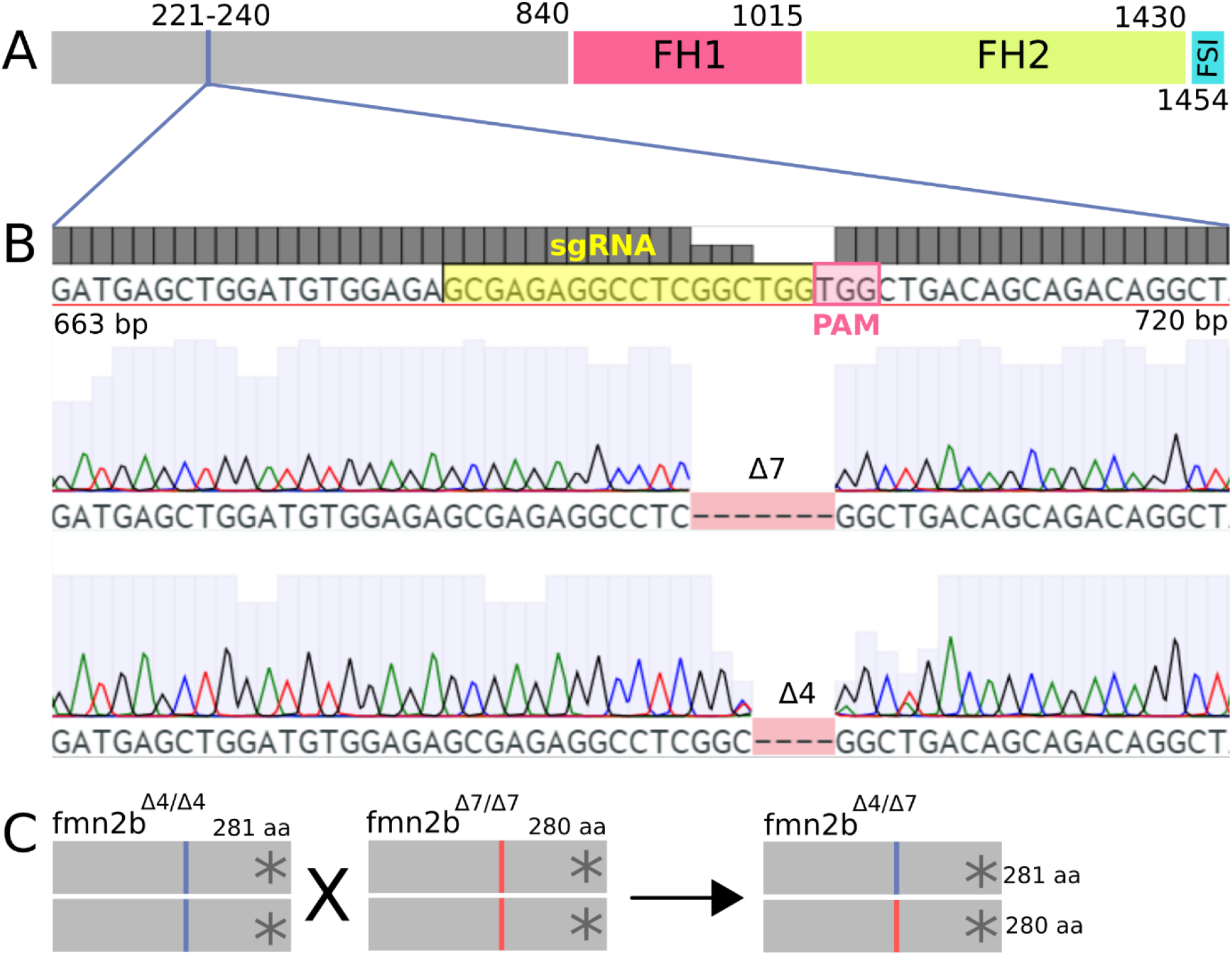
Generation of *fmn2b* CRISPR mutants. **A)** Schematic of the *fmn2b* protein indicating the functional protein domains and the site of *fmn2b* sgRNA-1. The FH1 domain spans the 841-1015 amino acids, FH2 domain spans the 1016-1430 amino acids and the FSI domain spans the 1431-1454 amino acids. **B)** Genomic locus from *fmn2b* exon 1 starting from 663 bp to 720 bp (corresponding to 221-240 amino acids). The sgRNA sequence is highlighted in yellow followed by the PAM sequence in pink. Representative chromatograms of amplicons sequenced from homozygous CRISPR mutants for *fmn2b* show two alleles with a 7 bp and a 4 bp deletion, respectively. The two alleles are denoted as *fmn2b*^Δ7^ (p.Leu233TrpfsTer281) and *fmn2b*^Δ4^ (p.Val234ThrfsTer280). **C)** Schematic outlining the generation of *fmn2b* homozygous mutants. Fish homozygous for two alleles are denoted as *fmn2b*^Δ7/Δ7^ and *fmn2b*^Δ4/Δ4^, the heteroallelic combination is denoted as *fmn2b*^Δ4/Δ7^.

### fmn2b mutants exhibit Spontaneous Tail Coiling (STC) and Touch-Evoked Escape Response (TEER) deficits

The first motor behaviour to appear in the developing zebrafish embryo is spontaneous tail coiling (STC) which starts around 17 hpf and persists till 27 hpf (Saint-Amant and Drapeau, 1998; Brustein et al., 2003). We recorded the spontaneous tail coiling in *fmn2b^+/+^* and *fmn2b^Δ4/Δ7^* embryos at 22 hpf (Movie 1). The frequency of spontaneous tail coiling was reduced in *fmn2b^Δ4/Δ7^* mutant embryos (3.168 ± 0.314 min^−1^) as compared to the wild-type *fmn2b^+/+^* embryos (5.003 ± 0.203 min^−1^) (Figure 2 A). However, the maximum amplitude of tail coiling did not change significantly in the *fmn2b^Δ4/Δ7^* embryos (Figure 2 B). The decrease of coiling frequency but not the magnitude of coiling indicates that the defects are likely due to deficits in the motor neuron or the neuromuscular junction (NMJ) function rather than loss of muscle integrity.

**Figure 2.**
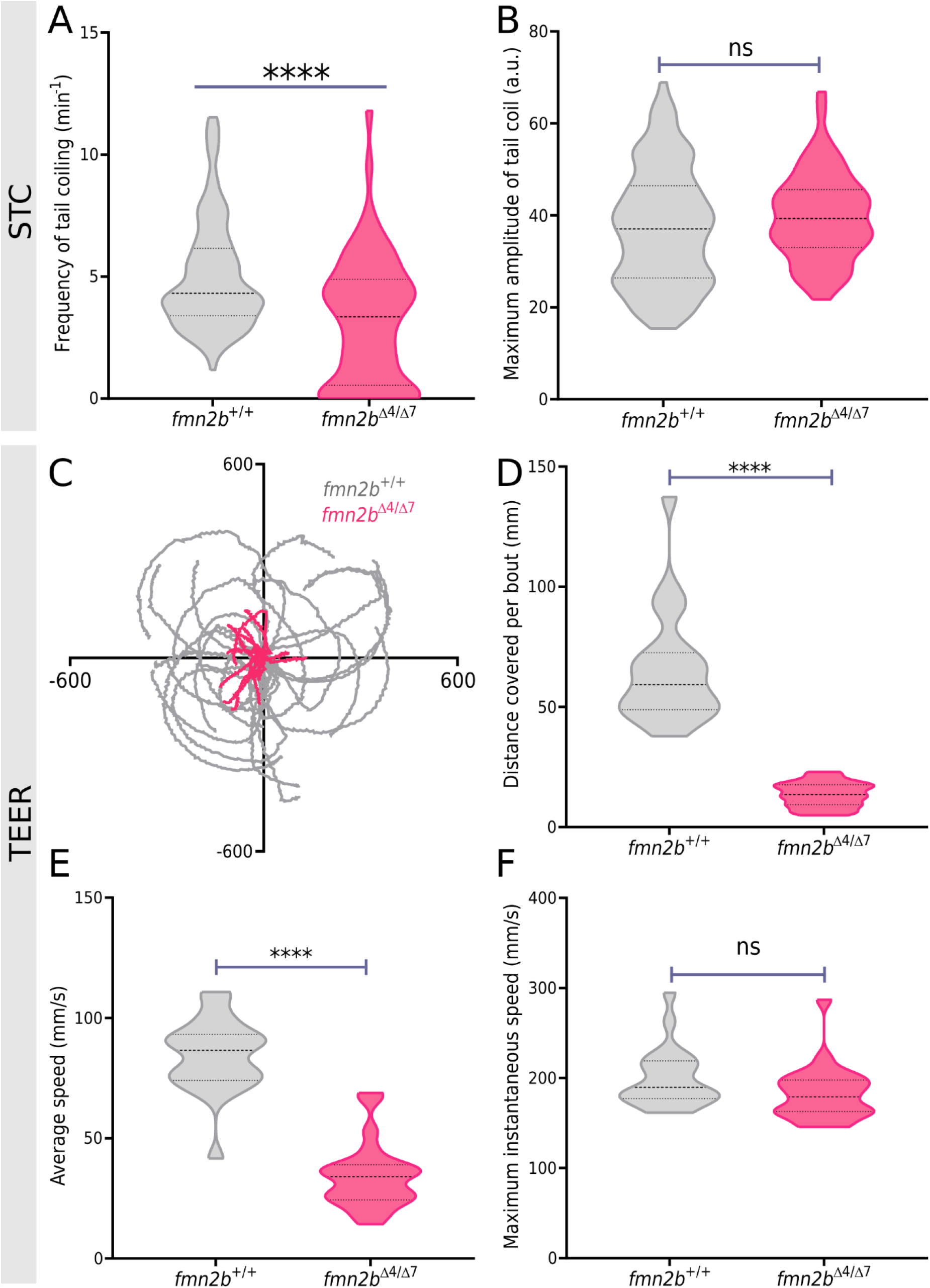
*fmn2b* mutants exhibit motor defects. **A)** Spontaneous tail coiling (STC) frequency and **B)** maximum amplitude of tail coiling in 22 hpf *fmn2b*^+/+^ (n=121) and *fmn2b^Δ4/Δ7^* (n=68) embryos. **C)** Trajectories of 2 dpf *fmn2b^+/+^* (grey traces; n=22) and *fmn2b^Δ4/Δ7^* (pink traces; n=24) embryos in Touch Evoked escape response (TEER), re-centered to a common origin for representation. The x and y axes in the plot correspond to the pixel values corresponding to coordinates of the fish from consecutively acquired frames. Quantification of **D)** distance covered per swim bout, **E)** average speed during the swim bout, and **F)** maximum instantaneous swimming speed of 2 dpf *fmn2b*^+/+^ (n=22) and *fmn2b*^Δ4/Δ7^ (n=24) embryos in response to a tactile stimulus. (**** p-value <0.0001; ns-not significant; Mann-Whitney U test)

As the embryo develops, the motor circuits mature to mediate touch-evoked escape responses (TEER) beginning at 48 hpf. TEER is typically manifested as a fast C-bend escape followed by a swimming bout. We performed the TEER assay on *fmn2b^+/+^* and *fmn2b^Δ4/Δ7^* embryos at 60 hpf (Movie 2) and recorded their response at 200 fps. Both the distance traversed (13.55 ± 1.06 mm) and the average speed (34.26 ± 2.785 mm/s) of touch-evoked swimming was reduced in 60 hpf *fmn2b^Δ4/Δ7^* embryos as compared to the distance covered (65.31 ± 4.93 mm) and swim speed (84.37 ± 3.253) of *fmn2b^+/+^* embryos (Figure 2 C, D, E). However, the maximum instantaneous speed attained during the entire swim bout was not significantly different between *fmn2b^+/+^* (199.5 ± 7.039 mm/s) and *fmn2b^Δ4/Δ7^* embryos (183.2 ± 6.245 mm/s) (Figure 2 F).

The comparable maximum tail coiling amplitudes and maximum instantaneous speeds observed in *fmn2b^+/+^* and *fmn2b^Δ4/Δ7^* embryos suggest abnormal motor neuron development or NMJ function and is unlikely to involve deficits in the musculature. Consistent with this, phalloidin staining of the *fmn2b^Δ4/Δ7^* embryos revealed no structural deformities in the axial muscles compared to the *fmn2b^+/+^* embryos (Figure S1).

### fmn2b mRNA is expressed in the spinal cord and spinal motor neurons of zebrafish embryos

The time window of STC and TEER in zebrafish embryos coincides with the outgrowth and pathfinding of primary motor neurons (Myers et al., 1986). *fmn2b* mRNA was found to be expressed in the spinal cord 48 hpf onwards suggesting a possible role in the development of spinal neurons (Figure 3 A, B). Previous reports have also reported Fmn2 mRNA expression in the spinal cord of chick, mouse and human embryos (Leader and Leder, 2000; Sahasrabudhe et al., 2016). To explicitly test the expression of *fmn2b* mRNA in the motor neurons, GFP positive motor neurons were isolated using Fluorescence-activated cell sorting (FACS) from 24 hpf and 60 hpf *Tg(mnx1:GFP)* embryos and the expression of *fmn2b* evaluated by RT-PCR. *fmn2b* transcript was expressed in motor neurons expressing GFP driven by the Tg(mnx1:GFP) isolated from both 24 hpf and 60 hpf embryos (Figure 3 C). Therefore, *fmn2b* is expressed in the spinal cord and in the spinal motor neurons of zebrafish embryos and could mediate motor neuron development and function.

**Figure 3.**
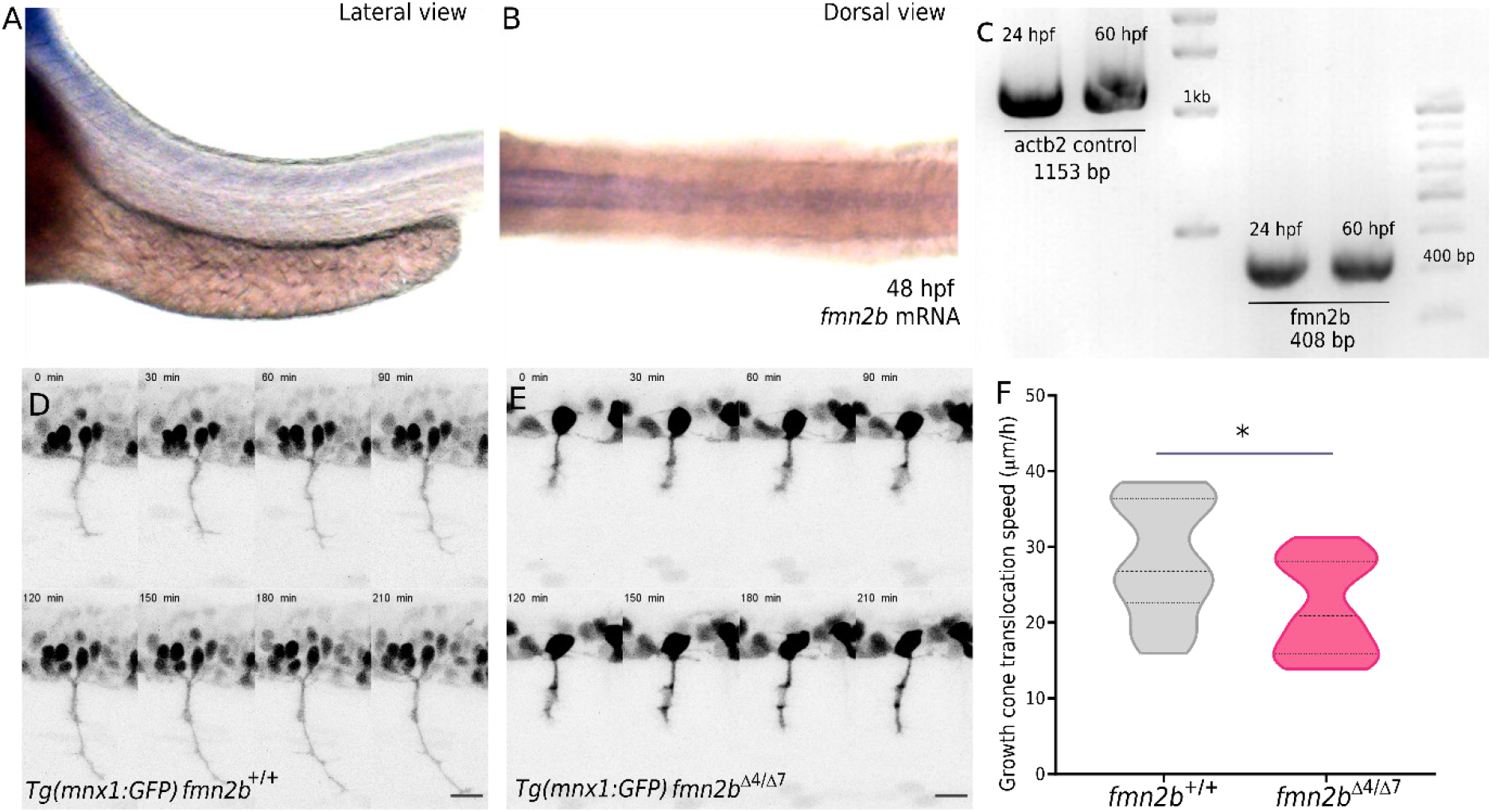
*fmn2b* is expressed in the spinal cord and motor neurons of zebrafish embryos. *fmn2b* is required for motor neuron growth cone translocation. **A)** Lateral and **B)** dorsal views of a representative 60 hpf embryo showing *fmn2b* mRNA expression in the spinal cord. **C)** Gel showing bands for amplified β2-actin (actb2) control and *fmn2b* from cDNA obtained from FACS sorted motor neurons (mnx1:GFP positive cells) isolated from 24 hpf and 60 hpf wild-type *Tg(mnx1:GFP)* embryos. **D)** Representative montages of live imaging of growth cone translocation in *fmn2b^+/+^* and E) *fmn2b^Δ4/Δ7^* embryos in the background of *Tg(mnx1:GFP)* during the time period of 22 hpf to 26 hpf. **F)** Quantification of motor neuron growth cone translocation speed in *fmn2b^+/+^* (n=15 growth cones; N=3 embryos) and *fmn2b^Δ4/Δ7^* (n=16 growth cones; N=4 embryos). (* p-value = 0.0298; Mann-Whitney U test). Scale bar is equivalent to 20 µm.

### CaP motor neuron outgrowth is slower in fmn2b mutants

To test the hypothesis that the motor defects in *fmn2b* mutants may arise due to deficits in motor neuron morphogenesis, the *fmn2b^Δ4/Δ7^* mutant line was crossed with the *Tg(mnx1:GFP)* line to obtain homozygous *fmn2b* mutants in the transgenic background of GFP-labelled motor neurons. The development of motor neurons was observed from 22 hpf to 26 hpf using time-lapse confocal imaging of the GFP-labelled neurons (Movie 3). The growth cones of the pioneering caudal primary (CaP) motor neurons were tracked using the Manual Tracking plugin in ImageJ, and the average translocation speed was calculated. The *fmn2b^Δ4/Δ7^* embryos showed slow outgrowth of the CaP neurons (Figure 3 D, E). The growth cone translocation speed of *fmn2b^Δ4/Δ7^* embryos (22.1 ± 1.575 µm/h) was slower than the *fmn2b^+/+^* embryos (28.34 ± 1.962 µm/h) (Figure 3 F).

### Primary motor neuron outgrowth and branching defects in fmn2b mutants

In addition to the slow outgrowth of CaP motor neurons in *fmn2b* mutants, collateral branches extended by the motor neurons at 24 hpf (Figure 4 A, B) and 60 hpf (Figure 4 G, H) were observed using the *fmn2b* mutant line *fmn2b^Δ4/Δ7^* in the background of the transgenic line *Tg(mnx1:GFP).* The embryos were mounted laterally to image the motor neurons and the images traced and quantified using the NeuronJ plugin in ImageJ. At 24 hpf, 100% of the hemisegments had the CaP motor neuron soma present in *fmn2b^+/+^* as well *fmn2b^Δ4/Δ7^* embryos. The branch density along the fascicle length (Figure 4 E) and the length of the motor fascicle (Figure 4 F) extended by the primary motor neurons at 24 hpf was found to be reduced in *fmn2b^Δ4/Δ7^* embryos. Similarly, 60 hpf *fmn2b^Δ4/Δ7^* embryos showed a reduction in the density (Figure 4 K) of collateral branches along the motor fascicle along with a reduction in the fascicle length (Figure 4 L). Similar defects were observed in homozygous *fmn2b* mutants with the *fmn2b^Δ4^* and *fmn2b^Δ7^* alleles (Figure S3).

**Figure 4.**
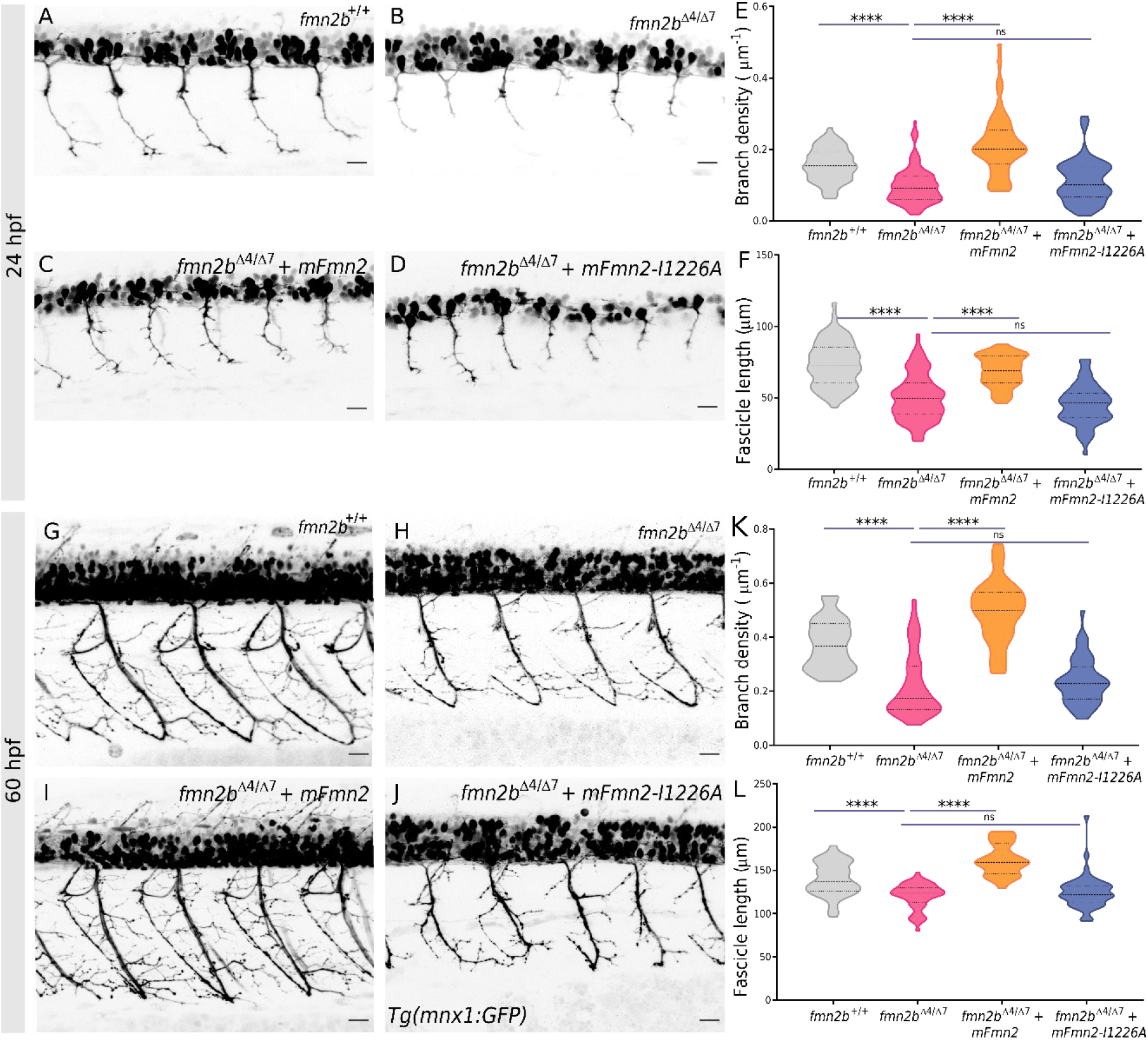
Motor neuron outgrowth and branching defects in *fmn2b* mutants. Representative micrographs of motor neurons labelled by *Tg(mnx1:GFP)* in 24 hpf **A)** *fmn2b*^+/+^ and **B)** *fmn2b*^Δ4/Δ7^ and 60 hpf **G)** *fmn2b*^+/+^ and **H)** *fmn2b*^Δ4/Δ7^ embryos. Motor neurons visualized by *Tg(mnx1:GFP)* in *fmn2b*^Δ4/Δ7^ mutant embryos injected at 1-cell stage with full-length mouse Fmn2 (mFmn2) mRNA at **C)** 24 hpf and **I)** 60 hpf and nucleation dead mouse Fmn2 (mFmn2-I1226A) mRNA at **D)** 24 hpf and **J)** 60 hpf. **E)** Quantification of branch density (number of branches per fascicle normalized to the fascicle length) per myotome hemisegment in 24 hpf *fmn2b*^+/+^ (0.1571 ± 0.005 µm^−1^; n=70 hemisegments) and *fmn2b*^Δ4/Δ7^ embryos (0.0981 ± 0.005 µm^−1^; n=118 hemisegments). The defects were rescued in *fmn2b*^Δ4/Δ7^ embryos injected with mFmn2 mRNA (0.2139 ± 0.012 µm^−1^; n=51 hemisegments) but not in the embryos injected with mFmn2-I1226A mRNA (0.111 ± 0.007 µm^−1^; n=63 hemisegments). **F)** Quantification of fascicle length extended by motor neurons per myotome hemisegment in 24 hpf *fmn2b*^+/+^ (68.97 ± 1.55 µm; n=70 hemisegments) and *fmn2b*^Δ4/Δ7^ embryos (44.98 ± 1.756 µm; n=118 hemisegments). The defects were rescued in *fmn2b*^Δ4/Δ7^ embryos injected with mFmn2 mRNA (73.7 ± 1.86 µm; n=51 hemisegments) but not in the embryos injected with mFmn2-I1226A mRNA (73.7 ± 1.86 µm; n=63 hemisegments). **K)** Quantification of branch density (number of branches per fascicle normalized to the fascicle length) per myotome hemisegment in 60 hpf *fmn2b*^+/+^ (0.3746 ± 0.014 µm^−1^; n=37 hemisegments) and *fmn2b*^Δ4/Δ7^ embryos (0.2211 ± 0.015 µm^−1^; n=57 hemisegments). The defects were rescued in *fmn2b*^Δ4/Δ7^ embryos injected with mFmn2 mRNA (0.4955 ± 0.021 µm^−1^; n=30 hemisegments) but not in the embryos injected with mFmn2-I1226A mRNA (0.2369 ± 0.012 µm^−1^; n=48 hemisegments). **L)** Quantification of fascicle length extended by motor neurons per myotome hemisegment in 60 hpf *fmn2b*^+/+^ (139.6 ± 3.21 µm; n=37 hemisegments) and *fmn2b*^Δ4/Δ7^ embryos (120.4 ± 1.94 µm; n=57 hemisegments). The defects were rescued in *fmn2b*^Δ4/Δ7^ embryos injected with mFmn2 mRNA (162.7 ± 3.38 µm; n=30 hemisegments) but not in the embryos injected with mFmn2-I1226A mRNA (124.7 ± 2.81 µm; n=48 hemisegments). (**** p-value <0.0001; ns - not significant; Kruskal Wallis test followed by Dunn’s post-hoc analysis). Scale bar is equivalent to 20 µm.

These observations implicate *fmn2b* function in both the outgrowth of the CaP motor neuron and collateral branching of motor neurons.

### F-actin nucleation activity of Fmn2 is required for motor neuron branching

The functional domains characteristic of Formin-2 are conserved across vertebrates and ectopic expression of full-length mouse Fmn2 (mFmn2) can rescue neurodevelopmental defects induced by morpholino-mediated knockdown of Fmn2b in zebrafish (Nagar et al., 2021). To assess if the expression of exogenous Fmn2 could rescue the outgrowth and branching defects, *mFmn2* mRNA tagged with mCherry was injected in the *fmn2b^Δ4/Δ7^* mutant embryos at the 1-cell stage. The branch density and fascicle length defects were rescued in both 24 hpf (Figure 4 C, E, F) and 60 hpf (Figure 4 I, K, L) mutant embryos.

F-actin nucleation is the canonical function of formins, including Fmn2, and is mediated by the FH2 domain (Goode and Eck, 2007; Quinlan et al., 2007; Yoo et al., 2015). The FH2 domain is conserved across phyla, including zebrafish (Nagar et al., 2021). A conserved isoleucine residue (at position 1226 in mouse Fmn2) is known to be critical for actin nucleation (Quinlan et al., 2007; Roth-Johnson et al., 2014; Kundu et al., 2020). Clustal omega alignment of the conserved isoleucine residue is shown in Figure S2. To test if actin nucleation by Fmn2b is required in zebrafish motor neuron morphogenesis, an F-actin nucleation dead version of mFmn2 with a point mutation converting the Isoleucine at 1226 amino acid position to Alanine (mFmn2-I1226A) was used. The mRNA for mFmn2-I1226A tagged with mCherry was injected in the *fmn2b^Δ4/Δ7^* embryos at the 1-cell stage and the motor neuron development was analyzed.

The *mFmn2-I1226A* mRNA injected 24 hpf (Figure 4 D, E, F) and 60 hpf (Figure 4 J, K, L) mutant embryos continue to exhibit the outgrowth and branching defects similar to the *fmn2b^Δ4/Δ7^* embryos. Rescue of phenotype exhibited by the *fmn2b^Δ4/Δ7^* mutants by full-length mFmn2 but not the nucleation dead version, mFmn2-I1226A points towards the conserved function of Fmn2 in motor neuron development and highlights the significance of F-actin nucleation by Fmn2 in outgrowth and branching of motor neurons.

### Rostral Primary (RoP) motor neuron development is compromised in fmn2b mutants

Despite slower outgrowth rates and reduced branching in *fmn2b^Δ4/Δ7^* embryos, the CaP fascicle eventually reaches the extremity of the ventral musculature in the zebrafish flank. On the contrary, another class of primary motor neurons, the Rostral primary (RoP) motor neurons which typically project to the mid-dorsal and mid-ventral musculature (Bagnall and McLean, 2014; Bello-Rojas et al., 2019), are permanently affected in *fmn2b^Δ4/Δ7^* embryos. The RoP soma are located at their predetermined site adjacent to the CaP and VaP cell body cluster in the spinal cord. The cell bodies of primary motor neurons are distinctly recognizable at 24 hpf.

On examining the 24 hpf embryos, the RoP cell body was found to be missing in *fmn2b* mutants. Compared to 2.5% of the hemisegments in *fmn2b^+/+^* embryos, 46.87% of the hemisegments in *fmn2b^Δ4/Δ7^* embryos did not have the RoP cell body. Injection of 1-cell stage *fmn2b^Δ4/Δ7^* embryos with *mFmn2* mRNA rescued this defect with only 2.85% hemisegments lacking RoP soma. However, the actin nucleation dead *mFmn2-I1226A* mRNA failed to rescue the loss of RoP soma in *fmn2b* mutants and 43.75% of the hemisegments continued to show loss of RoP soma (Figure 5 A-F).

**Figure 5.**
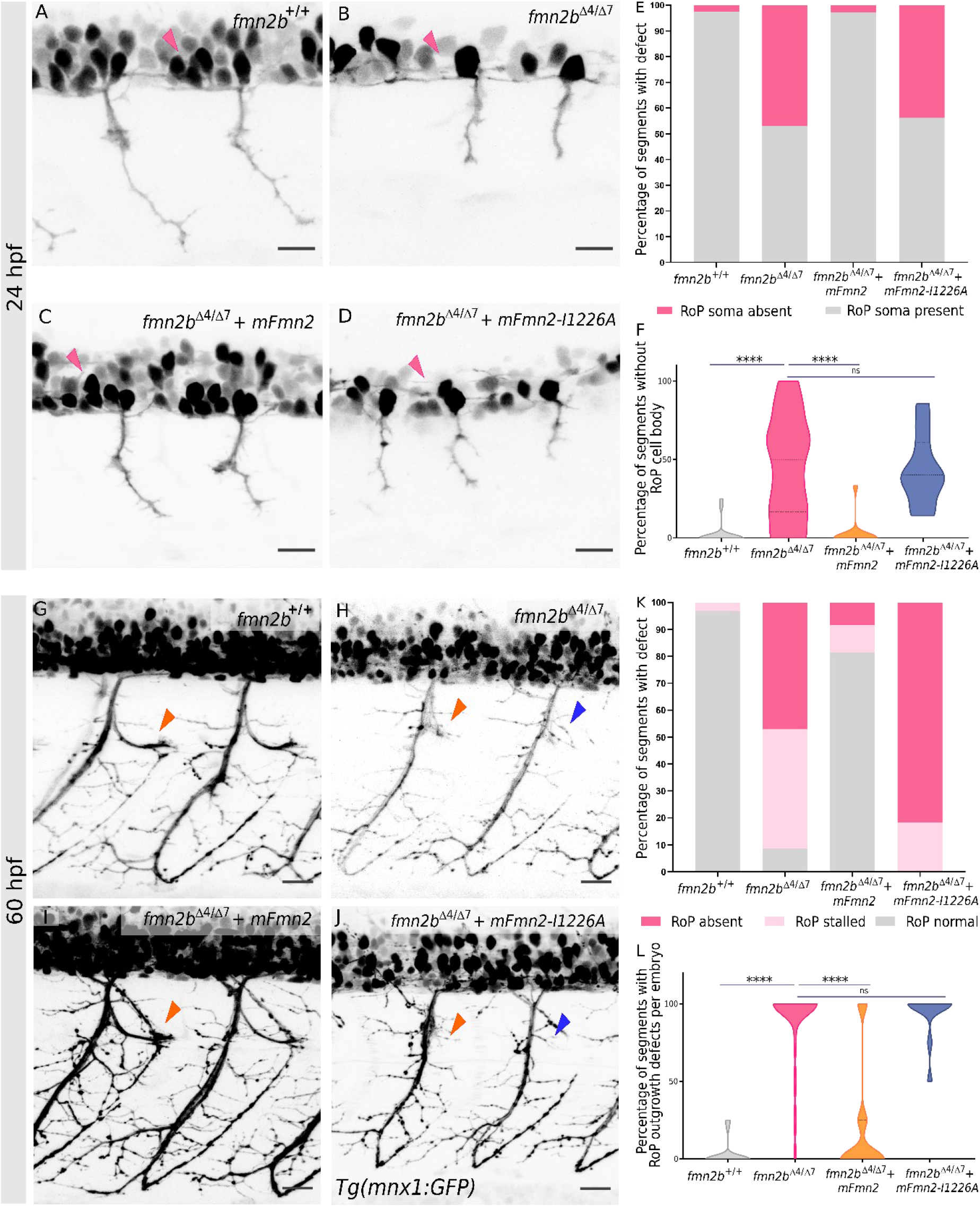
RoP soma and lateral innervation of the myotome by RoP are affected in *fmn2b* mutants. Representative micrographs of motor neurons labelled by *Tg(mnx1:GFP)* in 24 hpf **A)** *fmn2b*^+/+^ and **B)** *fmn2b*^Δ4/Δ7^ embryos. Representative micrographs of motor neurons labelled by *Tg(mnx1:GFP)* in 24 hpf old *fmn2b*^Δ4/Δ7^ mutant embryos injected at 1-cell stage with **C)** full-length mouse Fmn2 (mFmn2) mRNA and **D)** nucleation dead version, mFmn2-I1226A mRNA. The pink arrowheads indicate the RoP soma or its expected position. **E)** Bar graph summarizing the percentage of embryos with defects in RoP soma in 24 hpf *fmn2b*^+/+^ (n=18) and *fmn2b*^Δ4/Δ7^ embryos (n=23). The defects were rescued in *fmn2b*^Δ4/Δ7^ embryos injected with mFmn2 mRNA (n=12) but not in the embryos injected with mFmn2-I1226A mRNA (n=10). χ2 test of independence shows significant difference between the four groups, χ2 (df = 3) =101.8, p < 0.0001. **F)** Violin plots depicting the variation in data summarized in the bar graphs for 24 hpf embryos. (**** p-value <0.0001; ns - not significant; Kruskal Wallis test followed by Dunn’s post-hoc analysis). Scale bar is equivalent to 20 µm. Representative micrographs of motor neurons labelled by *Tg(mnx1:GFP)* in 60 hpf **G)** *fmn2b*^+/+^ and **H)** *fmn2b*^Δ4/Δ7^ embryos. Representative micrographs of motor neurons labelled by *Tg(mnx1:GFP)* in 60 hpf old *fmn2b*^Δ4/Δ7^ mutant embryos injected at 1-cell stage with **I)** full-length mouse Fmn2 (mFmn2) mRNA and **J)** nucleation dead version, mFmn2-I1226A mRNA. The orange arrowheads point towards the stalled RoP and the blue arrowheads indicate no RoP outgrowth. **K)** Bar graphs summarizing the percentage of embryos with defects in RoP axon outgrowth in 60 hpf *fmn2b*^+/+^ (n=22) and *fmn2b*^Δ4/Δ7^ embryos (n=26). The defects were rescued in *fmn2b*^Δ4/Δ7^ embryos injected with mFmn2 mRNA (n=14) but not in the embryos injected with mFmn2-I1226A mRNA (n=13). χ2 test of independence shows significant difference between the four groups, χ2 (df = 6) = 332.7, p < 0.0001. **L)** Violin plots depicting the variation in data summarized in the bar graphs for 60 hpf embryos. (**** p-value <0.0001; ns - not significant; Kruskal Wallis test followed by Dunn’s post-hoc analysis) Scale bar is equivalent to 20 µm.

At 60 hpf, the RoP motor neurons innervate their target in the mid-dorsal and mid-ventral musculature. The RoP motor neurons in *fmn2b^Δ4/Δ7^* mutants showed two major types of defects, absence of RoP outgrowth and stalling of RoP at the choice point near the horizontal myoseptum (HMS), where the RoP axons take a sharp lateral turn.

In the *fmn2b* mutant embryos, 91.3% of the hemisegments analyzed showed little to no lateral innervation. Of these, 44.35% of the hemisegments displayed defasciculated axons near the horizontal myoseptum, which acts as a choice point for the RoP axons to turn laterally. The hemisegments with defasciculated axons were classified as stalled RoP axons. Similar defects in RoP-like secondary motor neurons have been reported to occur due to depletion of Kif1b and Fidgetin like-1 in zebrafish (Fassier et al., 2018; Atkins et al., 2019). However, 46.95% of hemisegments had no RoP innervation altogether and were classified as RoP absent. Only 8.7% hemisegments showed the stereotypical RoP innervation and were classified as RoP present. In the *fmn2b^+/+^* embryos, only 3.15% of the hemisegments quantified showed RoP stalling defects, and the rest had stereotypical innervation by the RoP neurons. The RoP stalling and outgrowth defects were rescued by injection of full-length *mFmn2* mRNA with only 10.16% hemisegments with RoP stalling and 8.47% hemisegments with RoP outgrowth defects. On the other hand, RoP defects persisted in the *fmn2b^Δ4/Δ7^* embryos injected with *mFmn2-I1226A* mRNA, with 18.3% of hemisegments having stalled RoP axons and 81.7% without RoP innervation (Figure 5 G-L).

Collectively, these results indicate that *fmn2b* has pleiotropic effects on RoP development dependent on F-actin nucleation function. The deficits are at the level of loss of RoP soma, deficits in RoP axonogenesis and stalling at the horizontal myoseptum choice point.

### Overexpression of Fmn2 but not the nucleation dead mutant increases collateral branching in motor neurons

The role of *fmn2b* in motor neuron development has been established so far by observing *fmn2b* mutants, where the loss of Fmn2b in motor neurons causes a reduction in outgrowth and branching. Overexpression of mouse Fmn2 was employed in wildtype embryos to test the phenotypes in the context of Fmn2 gain of function. The embryos overexpressing mFmn2 showed increased branching, but overexpression of the nucleation dead mFmn2-I1226A did not cause any significant changes and was comparable to the wild-type embryos (Figure 6 A, B, E, F). The branch density was found to be increased in mFmn2 mRNA injected embryos at 24 hpf as well as 60 hpf but not in mFmn2-I1226A mRNA injected group (Figure 6 C, G). Interestingly, in 24 hpf embryos, the fascicle length was not significantly different in embryos overexpressing mFmn2 but was found to be decreased in embryos overexpressing mFmn2-I1226A. In 60 hpf embryos, the fascicle length was slightly increased in the mFmn2 overexpression group but remain unaffected in the mFmn2-I1226A overexpression group (Figure 6 D, H).

**Figure 6.**
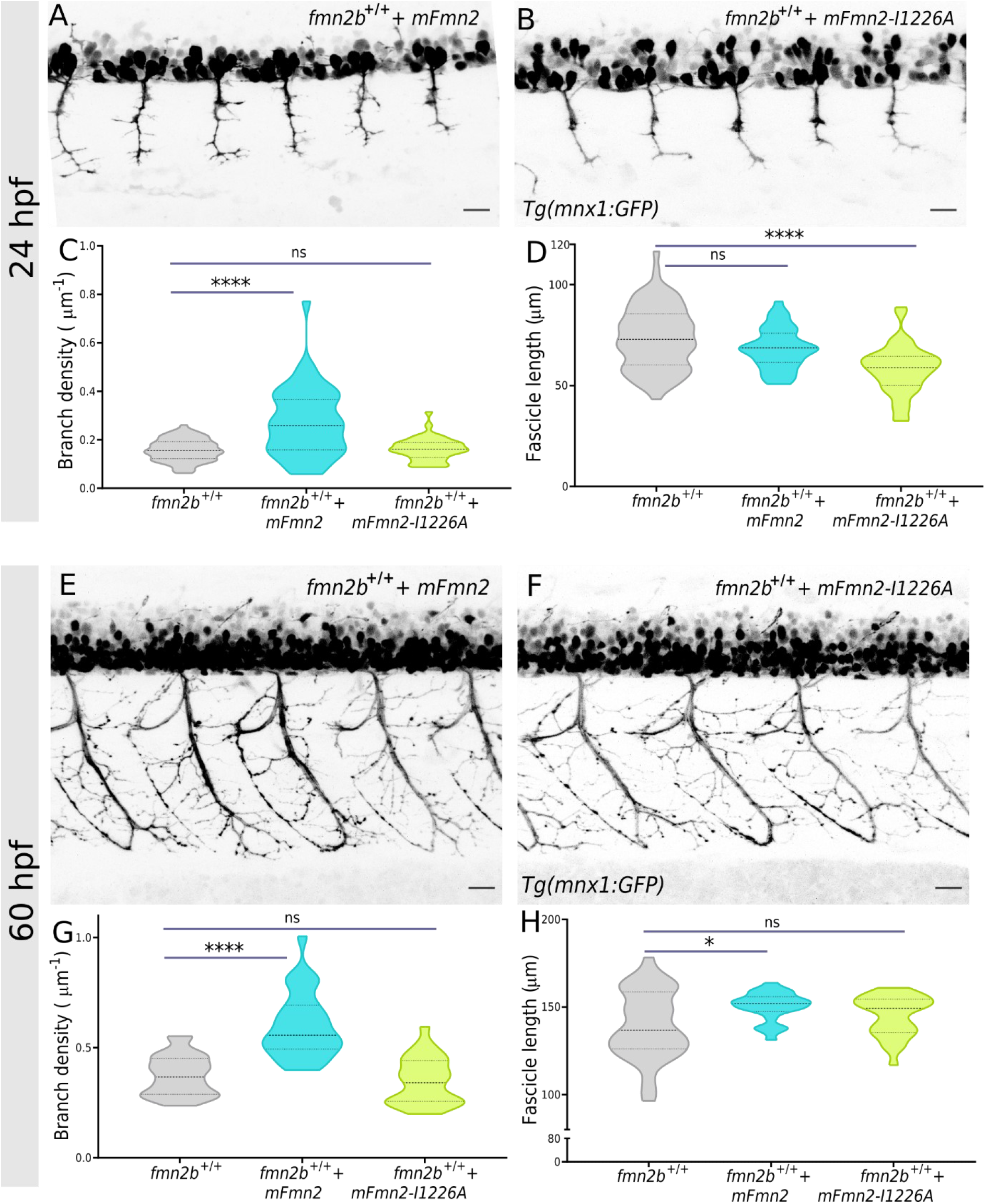
Outgrowth and branching phenotypes upon overexpression of mouse Fmn2 in wild-type embryos. Representative micrographs of motor neurons labelled by *Tg(mnx1:GFP)* in 24 hpf old *fmn2b*^+/+^ embryos injected with **A)** mFmn2 mRNA and **B)** mFmn2-I1226A mRNA. **C)** Quantification of branch density (number of branches per fascicle normalized to the fascicle length) along the fascicle extended by motor neurons per myotome hemisegment in *fmn2b*^+/+^ embryos injected with mFmn2 mRNA (0.2728 ± 0.0217 ^µm-1^; n=40 hemisegments) and mFmn2-I1226A mRNA (0.1617 ± 0.007 ^µm-1^; n=40 hemisegments). **D)** Quantification of fascicle length extended by motor neurons per myotome hemisegment in *fmn2b*^+/+^ embryos injected with mFmn2 mRNA (73.7 ± 1.86 µm; n=40 hemisegments) and mFmn2-I1226A mRNA (68.3 ± 1.603 µm; n=40 hemisegments). Representative micrographs of motor neurons labelled by *Tg(mnx1:GFP)* in 60 hpf old *fmn2b*^+/+^ embryos injected with **E)** mFmn2 mRNA and **F)** mFmn2-I1226A mRNA. **G)** Quantification of branch density (number of branches per fascicle normalized to the fascicle length) along the fascicle extended by motor neurons per myotome hemisegment in 60 hpf *fmn2b*^+/+^ embryos injected with mFmn2 mRNA (0.599 ± 0.027 ^µm-1^; n=28 hemisegments) and mFmn2-I1226A mRNA (0.3481 ± 0.02 ^µm-1^; n=28 hemisegments). **H)** Quantification of fascicle length extended by motor neurons per myotome hemisegment in 60 hpf *fmn2b*^+/+^ embryos injected with mFmn2 mRNA (150.4 ± 1.536 µm; n=28 hemisegments) and mFmn2-I1226A mRNA (145 ± 2.18 µm; n=28 hemisegments). (**** p-value <0.0001; * p-value = 0.0175; ns - not significant; Kruskal Wallis test followed by Dunn’s post-hoc analysis) Scale bar is equivalent to 20 µm.

However, quantification of RoP soma and RoP axon outgrowth phenotypes in the *fmn2b^+/+^* embryos overexpressing mFmn2 and mFmn2-I1226A did not reveal any significant changes (Figure S4).

### Neuromuscular Junction (NMJ) development in fmn2b mutants

Primary motor neurons innervate the fast-twitch muscles and form functional synapses by 48 hpf. To visualize the neuromuscular junctions (NMJ), double immunostaining of whole-mount 60 hpf embryos with znp-1 antibody (presynaptic marker) and α-bungarotoxin (post-synaptic marker) were undertaken. The co-localization of the pre- and post-synaptic markers revealed the engaged NMJ synapses in *fmn2b^+/+^* and *fmn2b^Δ4/Δ7^* embryos (Figure 7 A, B). Synapses (co-localization of znp-1 and α-bungarotoxin signals) along the neuronal arbours were quantified using the SynapCountJ plugin in Fiji. The number of synapses identified was normalized to the area of the myotome corresponding to one somite and the length of the neuronal arbour.

**Figure 7.**
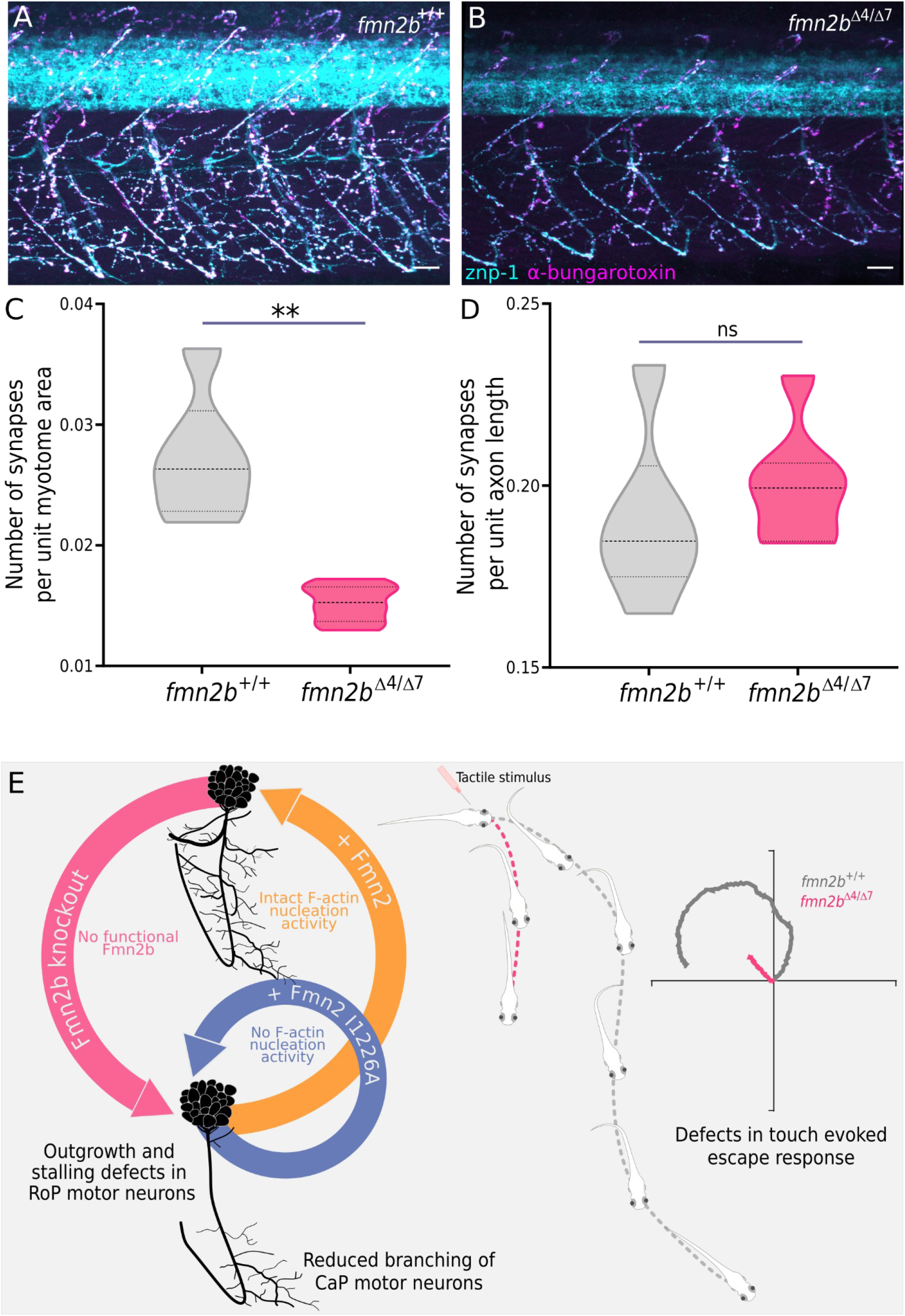
Synapse coverage in *fmn2b* mutants. Representative micrographs of 60 hpf **A)** *fmn2b*^+/+^ (n=6) **B)** *fmn2b*^Δ4/Δ7^ embryos (n=7) co-stained with znp-1 antibody and α-bungarotoxin. Quantification of colocalization of znp-1 and α-bungarotoxin in NMJ structures of 60 hpf embryos. **C)** Number of synapses per unit myotome area in *fmn2b*^+/+^ (0.0272 ±0.002; n=6) and *fmn2b*^Δ4/Δ7^ embryos (0.01521 ±0.0005; n=7, **p-value=0.0012). **D)** Number of synapses per unit axon length in *fmn2b*^+/+^ (0.1903± 0.009; n=6) and *fmn2b*^Δ4/Δ7^ embryos (0.2001± 0.005; n=7). (ns-not significant; Mann Whitney U test) Scale bar is equivalent to 20 µm. **E)** Depletion of Fmn2b causes outgrowth and branching defects in motor neurons in zebrafish larvae causing defects in motor behaviours. The outgrowth and branching defects in *fmn2b* mutants can be rescued by expression of full-length mouse Fmn2 in the mutants highlighting the requirement and conserved function of Fmn2 in development of motor neurons *in vivo*. The inability of mFmn2-I1226A to rescue the defects in *fmn2b* mutants underscores the requirement of the F-actin nucleation activity of Fmn2 in outgrowth and branching of motor neurons. The blue and orange curved arrows indicate the rescue experiments with full length mFmn2 and nucleation dead mFmn2-I1226A in the *fmn2b* mutant background respectively. The grey and pink traces indicate the trajectories of wildtype and *fmn2b* mutant embryos respectively in response to tactile stimulus. The schematic is not drawn to scale.

The *fmn2b^Δ4/Δ7^* mutants show reduced synaptic coverage of the myotomes compared to *fmn2b^+/+^* embryos (Figure 7 C). This result is consistent with previous observations of reduced outgrowth and branching density, and therefore the attenuated occupancy of myotome area, of motor neurons.

However, the number of synapses normalized to the arbour length was found to be comparable (Figure 7 D). Thus, while the ability of motor neurons to form synapses is intact in *fmn2b* mutants, the dramatically reduced arborization results in an effective reduction in the total number of synapses per myotome. The locomotor defects observed in *fmn2b* mutants is consistent with the decrease in the total number of synapses per myotome leading to insufficient activation of the myotome.

## DISCUSSION

Mutations, including loss-of-function mutations, in Fmn2 have been associated with intellectual disability, sensory dysfunction, age-associated cognitive decline (Perrone et al., 2012; Law et al., 2014; Agís-Balboa et al., 2017; Marco et al., 2018; Gorukmez et al., 2020). Interestingly, some affected individuals also develop hypotonia (Law et al). Further, mutations in Fmn2 have also been associated with sporadic amyotrophic lateral sclerosis in multiple studies (Schymick et al., 2007; Pamphlett et al., 2011; Morello et al., 2018). These associations suggest a possible involvement of Fmn2 in motor neuron development and function.

In this study, we generated *fmn2b* mutants using the CRISPR-Cas9 system (Figure 1) and found locomotor deficits in the homozygous mutants. *The fmn2b* mutants displayed deficits right from the earliest motor behaviours (STC at 22 hpf; Figure 2) to later stages where the development of the embryonic neural circuits is complete (TEER at 60 hpf; Figure 2). *fmn2b* mRNA expression was also detected in motor neurons and the spinal cord (Figure 3). Systematic evaluation of the primary spinal motor neurons revealed defects in the outgrowth of the CaP motor neurons. Live imaging indicated that the growth cone translocation velocity was reduced in *fmn2b^Δ4/Δ7^* embryos (Figure 3).

Recent *in vitro* studies in chick spinal neurons implicates Fmn2 in growth cone translocation. These studies implicate Fmn2 in mediating a molecular clutch that stabilizes the contact sites between the growth cone and the extracellular matrix and regulates the generation of traction forces required for outgrowth (Sahasrabudhe et al., 2016; Ghate et al., 2020). Additionally, Fmn2 facilitates the stabilization of protrusive processes, like growth cone filopodia, by coupling exploratory microtubules with the F-actin cytoskeleton (Kundu et al., 2021). Consistent with this, the depletion of Fmn2 reveals altered microtubule dynamics in the growth cones of zebrafish Rohon-Beard neurons *in vivo*. Similar mechanisms involving actin and microtubule remodelling could be involved in the translocation of CaP growth cones in zebrafish.

Even more striking than the delayed outgrowth rates of CaP motor neurons, the development of collateral branches which provide synaptic coverage of the entire muscle field was severely affected (Figure 4). The staggered development of spinal motor neurons in two phases allowed us to selectively look at the innervation of muscles by primary motor neurons at 24 hpf and collectively look at primary and secondary motor neuron innervation at 60 hpf. In addition, the 60 hpf embryos have elaborate branching and fully functional NMJ synapses allowing characterization of *fmn2b* in outgrowth, branching and synapse formation in motor neurons.

In 24 hpf *fmn2b* mutant embryos, branch density along the fascicle extended by the primary motor neurons and the fascicle length itself was reduced, suggesting a decrease in the innervation of the target fast muscle fibers (Figure 4). Similarly, the 60 hpf *fmn2b^Δ4/Δ7^* embryos have reduced branch density and fascicle outgrowth compared to wild-type embryos and are likely to result in reduced speed and distance covered by *fmn2b* mutants (Figure 4). Deficits in motor neuron outgrowth and collateral branching at both 24 hpf and 60 hpf was rescued by overexpressing mouse full length Fmn2 in the *fmn2b* homozygous mutants (Figure 4).

Axon collateral branching is a finely regulated multi-step process initiated by the rapid assembly of F-actin resulting in a filopodia-like protrusion from the axonal shaft (Ketschek and Gallo, 2010; Gallo, 2011, 2016; Coles and Bradke, 2015; Ketschek et al., 2015; Armijo-Weingart and Gallo, 2017; Menon and Gupton, 2018). Further cytoskeleton remodelling involving both the actin and microtubule cytoskeletons stabilize the protrusion and extend it to a collateral branch (Gallo, 2011, 2016; Armijo-Weingart and Gallo, 2017). The formation of the lateral protrusion from the axonal shaft is a major rate-limiting step in collateral branching and is initiated by the development of a juxtamembrane F-actin patch. *In vitro* studies implicate several actin binding proteins in regulating axonal F-actin patches. These include the actin nucleating Arp2/3 complex, Drebrin, WVE-1/WAVE regulatory complex (WRC) and cortactin (Spillane et al., 2011; Hu et al., 2012; Chia et al., 2014; Spillane and Gallo, 2014; Ketschek et al., 2016; Balasanyan et al., 2017). Recently, Fmn2 has been implicated in collateral branch formation in chick spinal neurons *in vitro*. Fmn2 appears to regulate the lifetime of axonal F-actin patches and consequently the probability of branch initiation (Kundu et al., 2020).

The F-actin nucleation and elongation activity of Fmn2 has been characterized *in vitro* (Montaville et al., 2014, 2016). Abrogation of F-actin nucleation activity of Fmn2 can be achieved by mutating a conserved Isoleucine residue in the FH2 domain to Alanine (Quinlan et al., 2007; Roth-Johnson et al., 2014; Kundu et al., 2020). We employed the full-length mouse Fmn2 bearing the isoleucine to alanine mutation (mFmn2-I1226A) in rescue experiments to test if the F-actin nucleating/elongating activity is necessary for motor neuron development.

Expression of mFmn2-I1226A in the *fmn2b* mutants could not rescue the outgrowth and branching defects of the motor neurons in 24 hpf and 60 hpf embryos (Figure 4). The failure of mFmn2-I1226A in rescuing the defects underscores the significance of the actin nucleating activity of Fmn2b in motor neuron outgrowth and branching consistent with reports from primary neuronal cultures of chick spinal cord reported previously (Kundu et al., 2020). Therefore, the motor neuron development in zebrafish is dependent on the F-actin nucleation activity of Fmn2b.

Loss of fmn2b had pleiotropic effects on the development of the RoP motor neuron. The RoP soma and RoP outgrowth were severely affected in *fmn2b* mutants. At 24 hpf, multiple hemisegments in *fmn2b* mutants did not have RoP soma at its characteristic position in the spinal cord (Figure 5). The role of Fmn2 in the regulation of differentiation or specification of progenitors is not formally tested despite several studies indirectly indicating pathways involving Fmn2 in cell differentiation. In a recent report, Fmn2 has been shown to cause neural progenitor differentiation defects Fmn2 and Flna double knockout mice in a synergistic manner (Lian et al., 2016). The absence of RoP soma in zebrafish *fmn2b* mutants may indicate a possible role of *fmn2b* in neural progenitor specification and/or differentiation. The RoP soma was seen in the expected location in *fmn2b^Δ4/Δ7^* embryos injected with mFmn2 mRNA but the defect could not be rescued by the injection of mFmn2-I1226A mRNA. Interestingly, the F-actin nucleating activity of Fmn2 is required for the differentiation of motor neuron progenitors or their specification. These results open up new possibilities to uncover the mechanistic role of Fmn2 in neural development.

In wild-type zebrafish embryos, the lateral projections from RoP motor neurons and follower secondary motor neurons begin outgrowth later than CaP and MiP neurons (Kuwada, 1993; Liu et al., 2016). In 60 hpf *fmn2b* mutants, the side branches of RoP motor neurons innervating the horizontal myoseptum were either not detected (46% embryos) or appeared to be stalled at the choice point (44% embryos), i.e., the horizontal myoseptum (Figure 5). RoP-like secondary motor neurons, which follow the same trajectory as RoP primary motor neurons have previously been shown to have pathfinding and stalling defects at the horizontal myoseptum in Fidgetin like-1 and Kif1b mutants (Fassier et al., 2018; Atkins et al., 2019). Similarly, the stalled RoP axons appear defasciculated in 60 hpf *fmn2b* mutants

One of the factors contributing to RoP outgrowth defects leading to the absence of RoP innervation in *fmn2b* mutants could be the lack of RoP soma as seen in 24 hpf mutant embryos. Further, the RoP stalling defect implicates Fmn2b in axonal pathfinding in response to guidance cues at the choice point. RoP outgrowth and stalling together caused a noticeable reduction in the innervation of the mid-dorsal and mid-ventral region of the axial myotome. The RoP may be more severely affected than the CaP neurons due to their late axonogenesis concomitant with the pleiotropic function and late expression of *fmn2b* in the spinal cord at 48 hpf.

Intriguingly, the overexpression of mouse Fmn2 in *fmn2b^+/+^* (wild-type) embryos causes opposite effects as compared to *fmn2b* knockout manifested as hyperbranching. This implies a significant role for *fmn2b* in regulating the collateral branching of motor neurons. Further, overexpression of the nucleation dead version of mFmn2 does not result in increased branch density and underscores the involvement of actin nucleation activity of Fmn2 (Figure 6).

Analysis of NMJ synapses using znp-1 and α-bungarotoxin double staining in *fmn2b* mutants showed no changes in the number of synapses along the total length of the motor neuron branches but showed a reduction in the total number of synapses when normalized to the area of the target myotome. We suggest that the Fmn2b has a primary role in regulating motor neuron branching and not in the formation of NMJ synapses. The behavioural defects in *fmn2b* mutants are likely due to the muscles not receiving sufficient input due to inadequate branching (Figure 7).

In a recent study, prolonged exposure of zebrafish larvae to strong and variable water currents caused upregulation of Fmn2b (Langebeck-Jensen et al., 2019). The rapid upregulation of Fmn2b in response to environmental stressors involving swimming and force generation in larvae with pre-established motor neural circuits invoke the possible involvement of *fmn2b* in neuronal plasticity and requires systematic investigation.

Taken together, the zebrafish model of Fmn2 loss-of-function offers unexpected insight into spinal motor neuron development and innervation of axial muscles. In addition to identifying the requirement of actin polymerization by Fmn2 in motor neuron morphogenesis and motor outputs, it highlights the central role of cytoskeleton remodelling in motor neuron homeostasis and the development of neuropathies.

## METHODS

### Zebrafish maintenance and procedures

All protocols used in this study were approved by the Institutional Animal Ethics Committee and the Institutional Biosafety Committee of IISER Pune or a National Human Genome Research Institute (NHGRI/NIH) Animal Care and Use Committee approved animal study protocol. The TAB5 wildtype strain of zebrafish was used for all the experiments including generation of CRISPR mutants. The TAB5 strain was used for all the wildtype outcrosses. Breeding pairs of adult zebrafish were maintained in recirculating aquaria (Techniplast) under a 14h-10h light-dark cycle. The temperature was maintained at 28.5°C and the pH was buffered between 7.2 to 7.8. The breeding adults were crossed to obtain embryos which were collected and grown in E3 buffer and used at different stages of development as indicated (Kimmel et al., 1995). For immunostaining and live imaging experiments, the buffer was supplemented with 0.003% Phenylthiourea (PTU; Sigma) to remove pigmentation from the skin. The transgenic line *Tg(mnx1:GFP)* was used to visualize motor neurons wherein the *mnx1* promoter specific to motor neurons drives GFP expression (Flanagan-Steet et al., 2005).

### Whole mount in situ hybridization

The RNA probes and procedure used for whole mount *in situ* hybridization experiments have been described previously (Nagar et al., 2021).

### Isolation of motor neurons from transgenic embryos, FACS, RT-PCR

Single cell suspension was made from around 200 *Tg(mnx1:GFP)* embryos as previously described (Bresciani et al., 2018). Briefly, the embryos were dissociated by trypsinization and filtering through a 70 µm sieve to obtain single cell suspension in 1X DMEM containing 10% FBS. The cells were sorted using a BD Biosciences fluorescence-activated cell sorting (FACS) equipment selecting cells expressing GFP, i.e., the motor neurons. RNA was extracted using Qiagen RNeasy Kit and cDNA was prepared using the SuperScript IV RT Kit (ThermoFisher). The cDNA was used for amplifying *fmn2b* transcripts in the mnx1 positive motor neurons. Primers and PCR protocol used to test the presence of *fmn2b* transcripts in motor neurons tagged by *Tg(mnx1:GFP)* were the same as the ones used for amplification of ISH probes from cDNA.

### RNA injections

Capped mRNA was synthesized using the HiScribe™ T7 ARCA mRNA Kit (with tailing) from linearized DNA template containing T7 promoter sequence upstream of the mFmn2-mCherry and mFmn2-I1226A-mCherry constructs. The transcribed mRNA was purified using RNeasy MinElute Cleanup Kit (Qiagen). 150 pg of mFmn2-mCherry and mFmn2-I1226A-mCherry mRNA was injected in the zebrafish embryos of desired genotype at 1-cell stage.

### sgRNA and Cas9 injections for CRISPR mutants

sgRNA targeting exon 1 of *fmn2b* was designed as previously described (Varshney et al., 2016). T7 HiScribe kit (NEB) was used to transcribe the sgRNA DNA template with the appended T7 promoter. The sgRNA was purified using ZymoResearch clean up columns. T3 mMessage mMachine RNA synthesis kit (Ambion) was used to synthesize capped Cas9 mRNA from the pT3TS-nCas9n plasmid (kind gift from Dr Wenbiao Chen; Addgene plasmid # 46757). The synthesized Cas9 mRNA was purified using the RNeasy MinElute Cleanup Kit (Qiagen). 30 pg sgRNA and 300 pg Cas9 mRNA was injected in zebrafish embryos at 1 cell stage. The sequence of sgRNA is given below.

Fmn2b_sgRNA : GGGCGAGAGGCCTCGGCTGG (ENSDARG00000061778.6; 12:47451436-47451459, plus strand)

### Genotyping CRISPR mutants

For identifying founder lines for *fmn2b* mutants, fluorescent PCR was performed and analyzed by capillary gel electrophoresis using the ABI GeneAnalyzer 3730XL, as previously described (Carrington et al., 2015; Varshney et al., 2015, 2016). Identification of homozygous mutant lines was also done using Sanger sequencing. Two zebrafish mutant lines with alleles causing a premature stop codon due to a frameshift mutation were established. The homozygous mutants were crossed to each other to obtain a heteroallelic mutant line for *fmn2b* to reduce the effect of any background mutations in the two mutant lines due to unintended off-target effects. Primer sequences for sanger sequencing of genomic DNA amplicons from crispants are as follows:

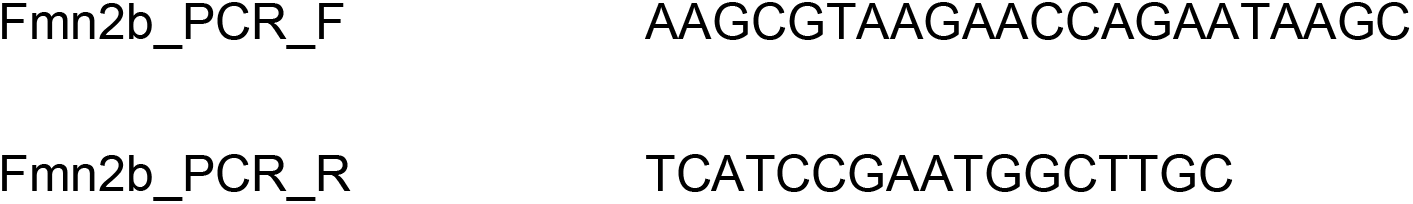

### Whole mount immunostaining

PTU treated embryos were collected at desired stages and fixed in 4% formaldehyde overnight at 4°C. For staining actin structures, fixed embryos were washed with 0.5% PBS-Triton, permeabilized with 2% PBS-Triton for 2 hours at room temperature and then incubated in Phalloidin Alexa Fluor 568 diluted 1:50 in 2% PBS Triton overnight at 4 °C in dark.

### Neuromuscular junction labelling and quantification

For neuromuscular junction staining of embryos, znp-1 antibody (DSHB; 1:100) and Tetramethyl Rhodamine labelled α-bungarotoxin (Invitrogen; 1:200) were used as pre-synaptic and post-synaptic markers respectively. Whole mount immunostaining procedures described in previous section were followed as described. The synapses were counted using the SynapCountJ plugin after making the traces of arbours in NeuronJ plugin in Fiji.

### Fluorescence microscopy, live imaging and mounting procedure

All the microscopic imaging was performed on the inverted LSM 780 confocal microscope (Zeiss) with a 25x oil immersion objective (NA 1.4). For imaging of the fixed samples, the embryos were cleared in 50% glycerol and mounted dorsal side down on a glass bottom petri dish using low gelling agarose (Sigma). For live imaging, the live embryos were mounted laterally in 0.5% low melt point agarose (Sigma) containing 0.003% MS-222 (Sigma) in a coverslip bottom 35 mm petri plate.

The growth cone of motor neurons was visualized using the *Tg(mnx1:GFP)* in wildtype or mutant background. The embryos were imaged starting at 22 hpf for 4-6 hours every 3 minutes. The growth cone translocation was analyzed using the Manual tracking plugin in Fiji to track the movement of the leading edge across time. The coordinates obtained were analyzed using the ibidi Chemotaxis tool to calculate the average growth cone translocation speed.

### Behaviour experiment set up and behaviour analysis

#### Spontaneous tail coiling (STC) assay

Embryos 22 hpf within their chorions were transferred to a 35 mm petri plate containing E3 buffer at of 28.5°C. A video camera (AVT Pike, F-032B) was used to record the spontaneous tail coiling behaviour of the embryos for a time period of 3 minutes at 15 fps. The videos were processed and analysed using a MATLAB script ZebraSTM published previously (González-Fraga et al., 2019).

#### Touch evoked escape response (TEER) assay

60 hpf zebrafish embryos were housed in a 35 mm petri plate containing pre-warmed E3 buffer at 28.5 °C. A tuberculin needle was repurposed by attaching a soft nylon fiber in front of the syringe holding the needle, to deliver tactile stimuli to zebrafish embryos. A soft touch was delivered to the head of the zebrafish once and their behaviour was recorded using a high-speed video camera (AVT Pike F-032B) at 208 fps. The videos obtained were analyzed using the Manual tracking plugin in Fiji to mark the trajectories of the zebrafish embryos upon receiving the tactile stimulus. The tracks were further analyzed using the ibidi Chemotaxis tool to calculate the distance travelled and average speed. The maximum instantaneous speed was calculated manually from the coordinates obtained from the manual tracking output.

### Figures and Statistical analysis

The analysis was performed for all the experiments in a genotype blinded manner. Image analysis was performed using Fiji and the Figure panels were assembled using Inkscape. The Violin plots were plotted in GraphPad Prism 8 and indicated statistical tests were performed using GraphPad Prism 8. The data is represented as mean ± SEM in the text. The statistical test used and the p values are indicated in the Figure legends.

## AVAILABILITY OF DATA AND MATERIALS

The data generated and analyzed in this study are included in this article and the additional information files. More information can be made available upon reasonable request to the corresponding author.

## ACKNOWLEDGEMENTS

The authors thank Dr. N. K. Subhedar (IISER Pune) for critical inputs on the manuscript. The authors acknowledge the IISER Pune Microscopy Facility, the National Facility for Gene Function in Health and Disease (NFGFHD) at IISER Pune and the zebrafish facility at NHGRI Zebrafish Core, NIH for access to equipment and infrastructure.

## AUTHOR INFORMATION

### AFFILIATIONS

1. **Indian Institute of Science Education and Research (IISER) Pune, Dr Homi Bhabha Road, Pune 411008, INDIA** Dhriti Nagar and Aurnab Ghose
2. **Zebrafish Core, Translational and Functional Genomics Branch, National Human Genome Research Institute, National Institutes of Health, Bethesda, Maryland, USA** Blake Carrington
3. **Translational and Functional Genomics Branch, National Human Genome Research Institute (NHGRI), National Institutes of Health (NIH), Bethesda, MD, USA** Shawn M Burgess

## CONTRIBUTION

Conceptualization: D.N. and A.G.; Investigation and formal analysis: D.N.; Methodology and resources: B.C. and S.M.B.; Writing – original draft: D.N. and A.G.; Writing – review and editing: D.N., B.C., S.M.B. and A.G; Funding Acquisition: A.G.. All authors gave final approval for publication and agreed to be held accountable for the work performed therein.

## CORRESPONDING AUTHORS

Correspondence: Dr. Aurnab Ghose, Indian Institute of Science Education and Research (IISER) Pune, Dr Homi Bhabha Road, Pune 411008, INDIA.

## ETHICS DECLARATIONS

### Ethics approval

All zebrafish husbandry and experimental protocols complied with institutional guidelines and were approved by the Institute Animal Ethics Committee (IAEC) and the Institutional Biosafety Committee (IBSC), IISER Pune or a National Human Genome Research Institute (NHGRI/NIH) Animal Care and Use Committee approved animal study protocol.

### Consent for publication

All authors gave consent for publication.

