## Supplementary Data for "Development of motor neurons and motor activity in zebrafish requires F-actin nucleation by Fmn2b"

### SUPPLEMENTARY FIGURES

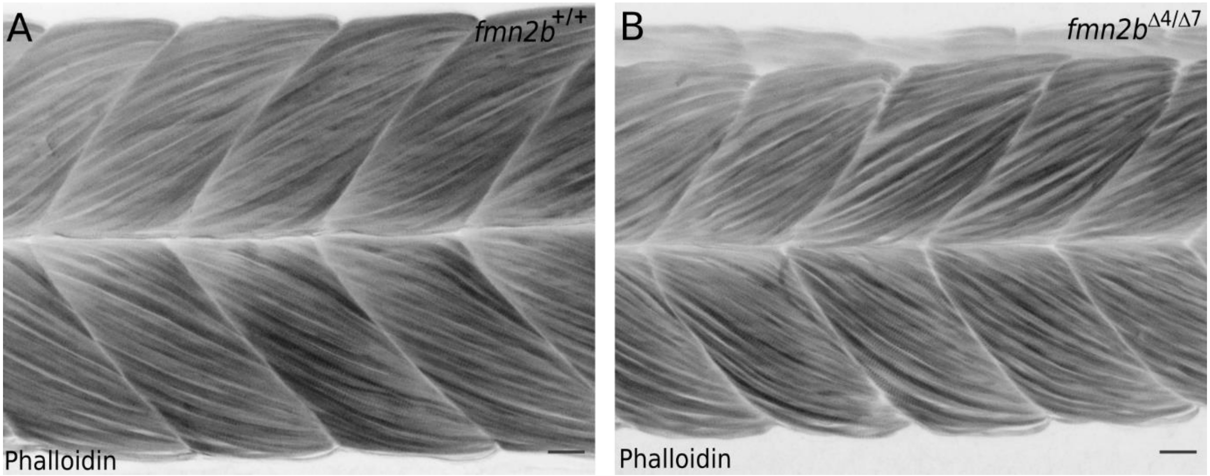


**Figure S1. Muscle integrity in *fmn2b* mutants**

Representative micrographs of 60 hpf **A)** *fmn2b*$^{+/+}$ (n=10) and **B)** *fmn2b*$^{\Delta 4/\Delta 7}$ (n=10) embryos stained with Alexa Fluor 568 tagged phalloidin to label the muscle structure. Scale bar is equivalent to 20 µm.

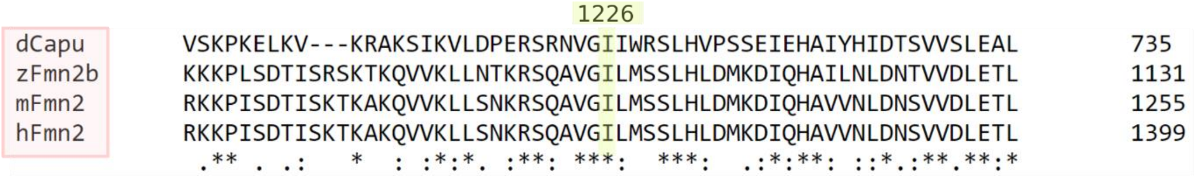


**Figure S2. I1226A residue is conserved across species**

Clustal Omega alignment of protein sequences flanking the conserved isoleucine residue responsible for F-actin nucleation activity in *Drosophila* Cappuccino (dCapu; Uniprot ID: Q24120), Zebrafish Fmn2b (zFmn2b; Uniprot ID: E7F517), mouse Fmn2 (mFmn2; Uniprot ID: Q9JL04) and human Fmn2 (hFmn2; Uniprot ID: Q9NZ56). The conserved isoleucine residue in mFmn2 is present at 1226 amino acid position highlighted in the alignment.

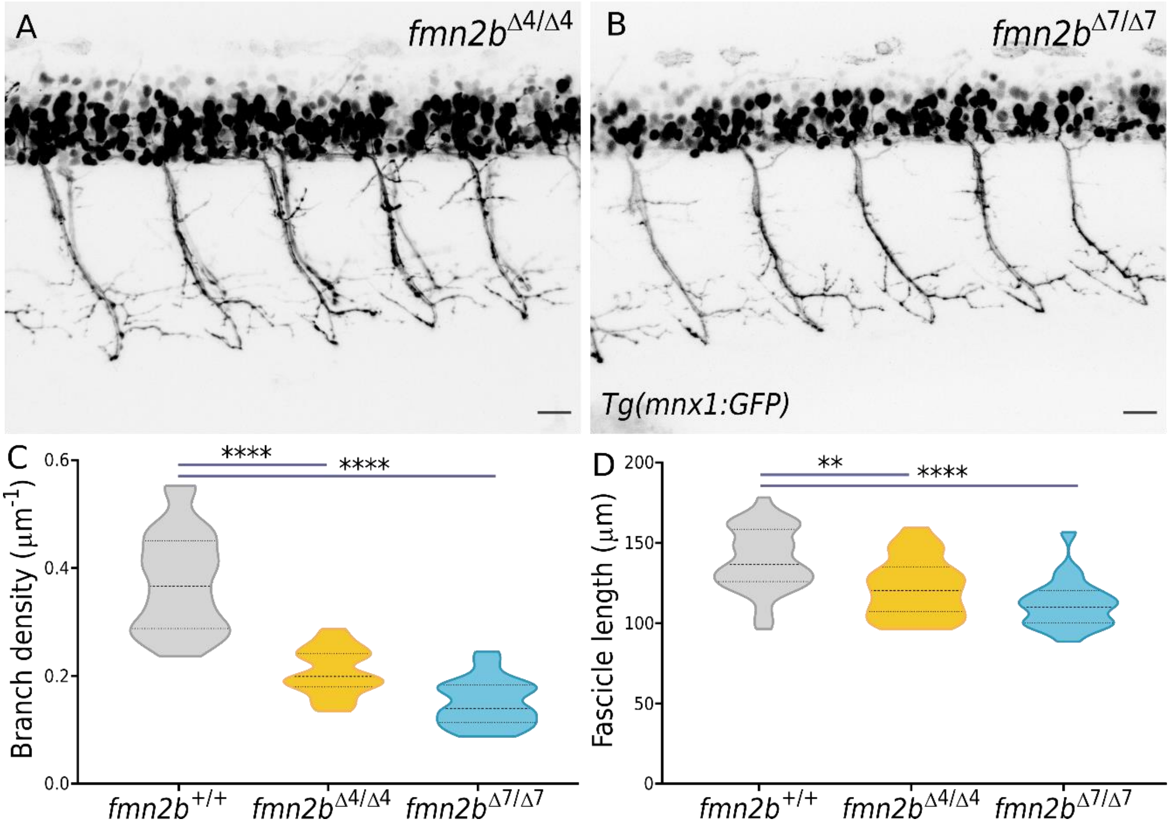


**Figure S3. Motor neuron branching defects in homoallelic *fmn2b* mutants**

The homozygous mutants with the *fmn2b*$^{\Delta7}$ and *fmn2b*$^{\Delta4}$ alleles also show motor neuron branching defects as observed in the *fmn2b*$^{\Delta4/\Delta7}$ embryos. Representative micrographs of motor neurons labelled by *Tg(mnx1:GFP)* in 60 hpf **A)** *fmn2b*$^{\Delta4/\Delta4}$ and **B)** *fmn2b*$^{\Delta7/\Delta7}$ embryos. **C)** Quantification of branch per myotome hemisegment in 60 hpf *fmn2b*$^{+/+}$ (0.3746 ± 0.014 $\mu m^{-1}$; n=37 hemisegments), *fmn2b*$^{\Delta4/\Delta4}$ embryos (0.2073 ± 0.009 $\mu m^{-1}$; n=22 hemisegments) and *fmn2b*$^{\Delta7/\Delta7}$ embryos (0.151 ± 0.009 $\mu m^{-1}$; n=21 hemisegments). **D)** Quantification of fascicle length extended by motor neurons per myotome hemisegment in 60 hpf *fmn2b*$^{+/+}$ (139.6 ± 3.21 µm; n=37 hemisegments), *fmn2b*$^{\Delta4/\Delta4}$ embryos (122.3 ± 3.93 µm; n=22 hemisegments) and *fmn2b*$^{\Delta7/\Delta7}$ embryos (111.9 ± 3.33 µm; n=21 hemisegments). (**** p-value <0.0001; ** p-value = 0.0065; ns - not significant; Kruskal Wallis test followed by Dunn's post-hoc analysis). Scale bar is equivalent to 20 µm.

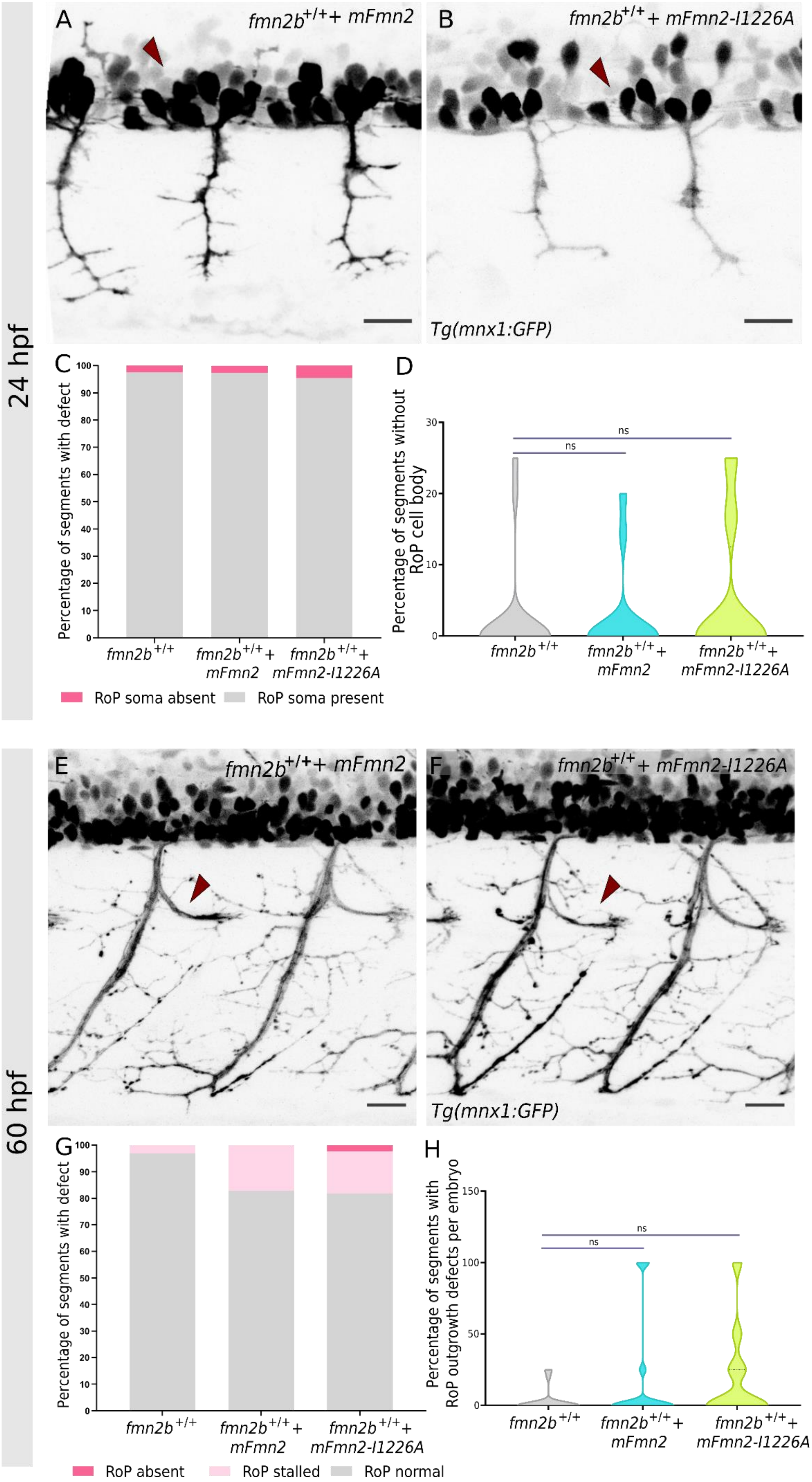
24 hpf
A
fmn2b+/++ mFmn2
B
fmn2b+/++ mFmn2-I1226A
Tg(mnx1:GFP)
C
Percentage of segments with defect
fmn2b+/+
fmn2b+/++ mFmn2
fmn2b+/++ mFmn2-I1226A
RoP soma absent
RoP soma present
D
Percentage of segments without RoP cell body
ns
ns
60 hpf
E
fmn2b+/++ mFmn2
F
fmn2b+/++ mFmn2-I1226A
Tg(mnx1:GFP)
G
Percentage of segments with defect
RoP absent
RoP stalled
RoP normal
H
Percentage of segments with RoP outgrowth defects per embryo

**Figure S4. RoP development is not affected by overexpression of Fmn2 in wild-type embryos**

Representative micrographs of motor neurons labelled by *Tg(mnx1:GFP)* in 24 hpf old *fmn2b*$^{+/+}$ embryos injected with **A)** mFmn2 mRNA and **B)** mFmn2-I1226A mRNA. Red arrowhead points towards the RoP soma. Representative micrographs of motor neurons labelled by *Tg(mnx1:GFP)* in 60 hpf old *fmn2b*$^{+/+}$ embryos injected with **E)** mFmn2 mRNA and F**)** mFmn2-I1226A mRNA. Red arrowhead points towards the RoP outgrowth. **C)** Bar graphs summarizing the percentage of embryos with defects in RoP cell body in 24 hpf *fmn2b*$^{+/+}$ (n=18). No defects were observed in *fmn2b*$^{+/+}$ embryos injected with mFmn2 mRNA (n=13) and mFmn2-I1226A mRNA (n=16). **G)** Bar graphs summarizing the percentage of embryos with defects in RoP axon outgrowth in 60 hpf *fmn2b*$^{+/+}$ (n=22). No defects were observed in *fmn2b*$^{+/+}$ embryos injected with mFmn2 mRNA (n=13) and mFmn2-I1226A mRNA (n=11). Violin plots depicting the variation in data summarized in the bar graphs for **D)** 24 hpf embryos and **H)** 60 hpf embryos. (ns- not significant; Kruskal Wallis test followed by Dunn's post-hoc analysis). Scale bar is equivalent to 20 µm.

**Movie legends**

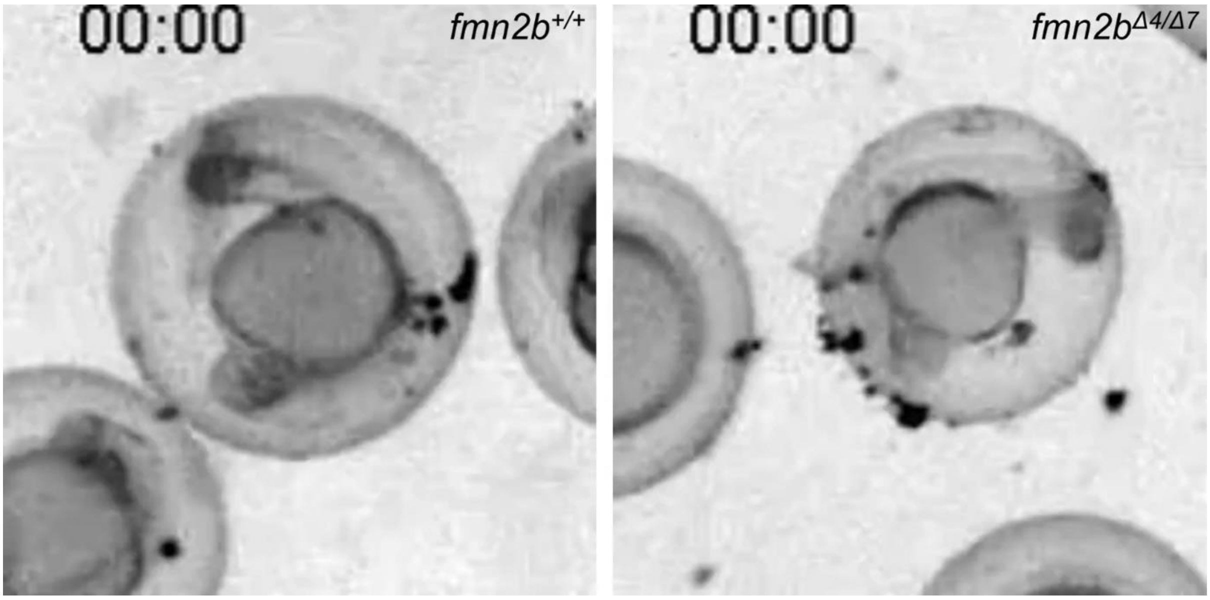


**Movie 1. Spontaneous tail coiling (STC) in 22 hpf *fmn2b*$^{+/+}$ and *fmn2b*$^{\Delta 4/\Delta 7}$ embryos.**

Spontaneous tail coiling response in 22 hpf *fmn2b*$^{+/+}$ (wildtype) and *fmn2b*$^{\Delta 4/\Delta 7}$ (*fmn2b* mutant) zebrafish embryos was recorded at 15 fps for a duration of 3.5 minutes.

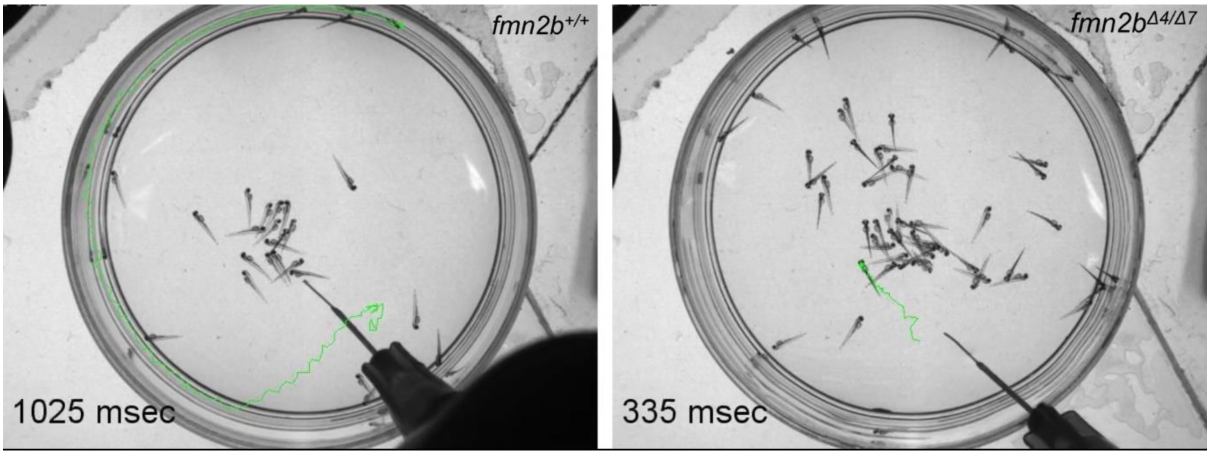


**Movie 2. Touch evoked escape response (TEER) in 60 hpf *fmn2b*$^{+/+}$ and *fmn2b*$^{\Delta 4/\Delta 7}$ embryos.**

Response of 60 hpf *fmn2b*$^{+/+}$ (wildtype) and *fmn2b*$^{\Delta 4/\Delta 7}$ (*fmn2b* mutant) zebrafish embryos to tactile stimulus was recorded at 200 fps for the duration of the swimming bout.

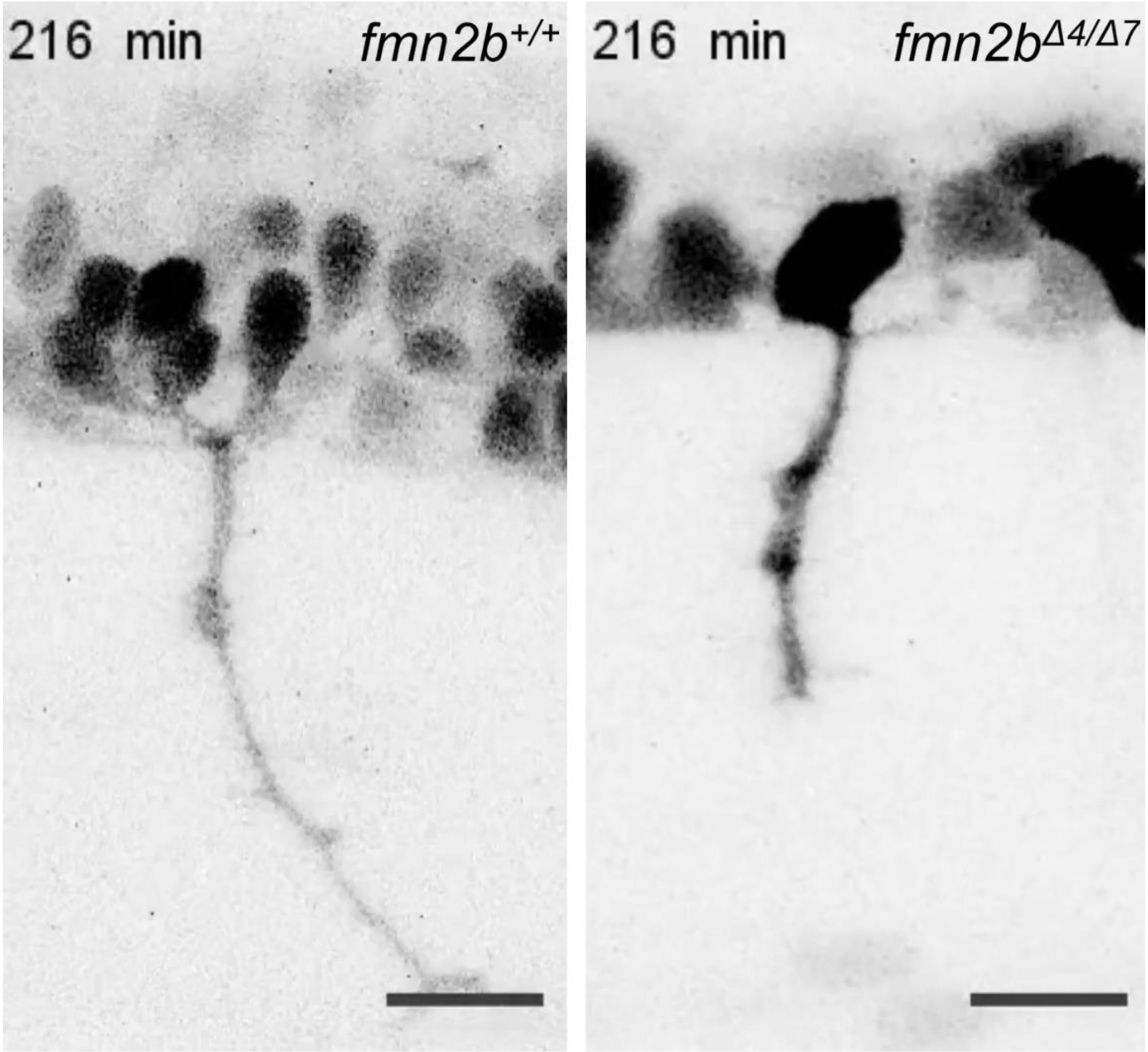


**Movie 3. CaP growth cone translocation in 18 hpf *fmn2b*$^{+/+}$ and *fmn2b*$^{\Delta 4/\Delta 7}$ embryos.**

Live imaging of caudal primary (CaP) motor neuron growth cone translocation visualized using the *Tg(mnx1:GFP)* during 22 hpf to 26 hpf using time-lapse confocal imaging. The images were acquired at a time interval of 3 minutes. Scale bar is equivalent to 20 µm.
